# Single-cell RNA-Seq analysis reveals dual sensing of HIV-1 in blood Axl^+^ dendritic cells

**DOI:** 10.1101/2022.01.03.474732

**Authors:** Flavien Brouiller, Francesca Nadalin, Ouardia Aït Mohamed, Pierre-Emmanuel Bonté, Constance Delaugerre, Jean-Daniel Lelièvre, Florent Ginhoux, Nicolas Ruffin, Philippe Benaroch

## Abstract

Sensing of incoming viruses represents one of the pivotal tasks of dendritic cells (DC). Human primary blood DC encompass various subsets that are diverse in their susceptibility and response to HIV-1. The recent identification of Axl^+^DC, a new blood DC subset, endowed with unique capacities to bind, replicate, and transmit HIV-1 prompted us to evaluate its anti-viral response. We show that HIV-1 induced two main broad and intense transcriptional programs in different Axl^+^DC potentially induced by different sensors; a NF-κB-mediated program that led to DC maturation and efficient antigen-specific CD4^+^T cell activation, and a program mediated by STAT1/2 that activated type I IFN and an ISG response. These responses were absent from cDC2 exposed to HIV-1 except when viral replication occurred. Finally, Axl^+^DC actively replicating HIV-1 identified by quantification of viral transcripts exhibited a mixed NF-κB/ISG innate response. Our results suggest that the route of HIV-1 entry may dictate different innate sensing pathway by DC.

## INTRODUCTION

Dendritic cells (DCs) are crucial for the recognition of pathogens and the initiation of adaptive immune responses (Schlitzer et al., 2015). In addition, DCs are thought to be important for the establishment and the dissemination of HIV-1 in the host and thus play both a role in antiviral immunity and HIV-1 transmission (Martín-Moreno and Muñoz-Fernández, 2019). Yet, how the various primary human DC detect and respond to HIV-1 remains poorly understood. While most of the studies of HIV-1 innate sensing by DC have been performed using *in vitro* differentiated DC (Yin et al., 2020), only a few studies have been performed with fresh primary human DCs (Beignon et al., 2005; Fonteneau Jean-François et al., 2004; Gringhuis et al., 2010; Silvin et al., 2017). Early studies on monocyte-derived DC (MDDC) demonstrated that HIV-1 retrotranscription (RT) is necessary to induce cell maturation, as jugged by expression of the costimulatory molecule CD86 (Manel et al., 2010). However, HIV-1 replication in myeloid cells is highly restricted by the SAMHD1 enzyme that degrades the cytosolic pool of dNTP, but is counteracted by HIV-2 coded protein Vpx (Cribier et al., 2013; Hrecka et al., 2011; Laguette et al., 2011; Lahouassa et al., 2012). Insight in the signalling pathways involved upon MDDCs exposure to HIV-1 in the presence of Vpx, led to the identification of the cGAS – STING pathway in the induction of an interferon response and the maturation of DC (Johnson et al., 2018; Lahaye et al., 2013; Manel et al., 2010). Whether similar pathways are activated in primary fresh DC populations remains to be determined.

Single-cell techniques have been pivotal to identify DC populations at high resolution, to establish new molecular cell markers, and to possibly infer their ontogeny (See et al., 2017; Villani et al., 2017). These studies identified a new human DC population expressing Axl (herein referred as Axl^+^DC). It shares markers with both plasmacytoid and conventional DC populations, but little is known about its role and functions (Alcántara-Hernández et al., 2017; Leylek et al., 2019; See et al., 2017; Villani et al., 2017). We have previously established that Siglec-1, one of the few specific Axl^+^DC markers, efficiently mediates HIV-1 capture and infection in Axl^+^DC (Ruffin et al., 2019). Incorporation of the HIV-2 Vpx accessory protein into HIV-1 particles highly increases the rates of Axl^+^DC infection (Ruffin et al., 2019). Infected Axl^+^DC were efficient at transmitting newly formed HIV-1 infectious particles to activated CD4^+^ T lymphocytes. Of note, HIV-1 assembly in Axl^+^DC occurs in apparently internal compartments previously observed in macrophages, while the assembly process in cDC2 occurs at the plasma membrane in a polarized manner similar to what is observed in infected CD4^+^T lymphocytes. In addition, whereas Axl^+^DC activation through Toll-like receptor (TLR) inhibits their infection by preventing viral fusion, activated Axl^+^DC still support HIV-1 transmission to CD4^+^ T lymphocytes via Siglec-1 (Ruffin et al., 2019), in a process called *trans*infection (Izquierdo-Useros et al., 2014). Axl^+^DC may thus participate to HIV-1 infection and spreading into the host.

Among the other blood DC populations, cDC2 (CD1c^+^DC) are also susceptible to HIV-1 infection, while cDC1 (CD141^+^DC) and plasmacytoid DC (pDC) are resistant (Ruffin et al., 2019; Silvin et al., 2017). When exposed to HIV-1, pDC develop a TLR7-mediated strong type I IFN response (Beignon et al., 2005; Silvin et al., 2017), while cDC2 require Vpx and viral replication to exhibit a small increased expression of CD86 and a production of IP-10 (Silvin et al., 2017). cDC1 response to HIV-1 in the presence of Vpx is independent of viral replication and remains lower in intensity than the one of cDC2 suggesting a link between viral replication and the response observed in cDC2 (Silvin et al., 2017). Moreover, innate sensing in cDC2 as well as in MDDC occurs via the cGAS-STING pathway which detects viral cDNA (Lahaye et al., 2018; Silvin et al., 2017).

Given the complex role of DC populations in HIV-1 infection which can favor the initial phases of the infection, but also can elicit anti-viral innate and adaptative responses (Rhodes et al., 2019; Sáez-Cirión and Manel, 2018), it was of interest to evaluate the Axl^+^DC innate response to incoming HIV-1 particles which very efficiently fuse with these cells (Ruffin et al., 2019). Here, we investigated the transcriptional response of *ex vivo* isolated Axl^+^DC upon HIV-1 exposure, the sensing pathways involved, and the impact on Axl^+^DC capacity to activate CD4^+^ T lymphocytes. We discovered that sensing of HIV-1 is set up by Axl^+^DC ahead of viral replication and that two independent responses are triggered in different cells. The main response of Axl^+^DC exposed to HIV-1 includes the activation of the STAT and IRF transcription factors, responsible for ISG expression. The second response is characterised by the activation of NF-κB transcription factor that leads to the maturation of Axl^+^DC. Such responses were characterized computationally using complementary approaches and validated *in vitro.*

## RESULTS

### Among the blood DC subsets, Axl^+^DC exhibit the stronger and wider innate response to HIV-1 exposure

To better characterize the interplay between HIV-1 and primary DC that are susceptible to viral infection, we investigated the transcriptional responses of sorted cDC2 and Axl^+^DC following HIV-1 (NL-AD8) exposure by bulk RNA sequencing (RNAseq) (Figure 1A). While levels of HIV-1 transcripts were similar in the two subsets (Figure 1B), changes in gene expression were more pronounced and more numerous in Axl^+^DC compared to cDC2 (Figure 1A and 1C). The presence of Vpx into the viral particles induced higher viral replication, particularly in Axl^+^DC, and led to bigger changes in gene expression in both cell types (Figure 1C and S1A-B). Inflammatory cytokines and interferon-stimulated genes (ISGs) were impacted in both cDC2 and Axl^+^DC by HIV-1 in the presence of Vpx. In contrast, genes of the NF-κB pathway (*CD40, CD80, CD86, NFKB,* and *REL*) were only induced in Axl^+^DC upon HIV-1(AD8) exposure, in the presence or absence of Vpx, and even in the presence of azidothymidine (AZT) and neverapin (NVP), which both inhibit retroviral transcription (Figure 1A and S1C). We confirmed at the protein level the preferential activation of ISG and inflammatory pathways of Axl^+^DC compared to cDC2 by measuring the expression of MX1/IP-10/CCL13 and CD86/IL-8 respectively, by flow cytometry (Figure S1D and S1E). IFN responses and TNFα signalling were induced by HIV-1 only in Axl^+^DC and independently of viral retrotranscription or replication (GSEA on differentially expressed genes (DEGs), Figure 1D and 1E). HIV-1-exposed Axl^+^DC up-regulate genes involved in DNA cytosolic sensing (that include ISGs), as well as NF-kB and TNF signalling (KEGG pathway enrichment analysis on DEGs, Figure 1F). In addition, we found activation of signalling pathways involving the cytosolic RNA sensor RIG-1 (retinoic acid-induced gene I), and Toll-like receptors (TLR) in Axl^+^DC (Figure 1F and S1F). Thus, these results indicate that Axl^+^DC are unique among DC subsets in the magnitude and amplitude of their response to HIV-1 exposure. This response includes activation of various sensors and signalling pathways, even prior to HIV-1 retrotranscription.

**Figure1:**
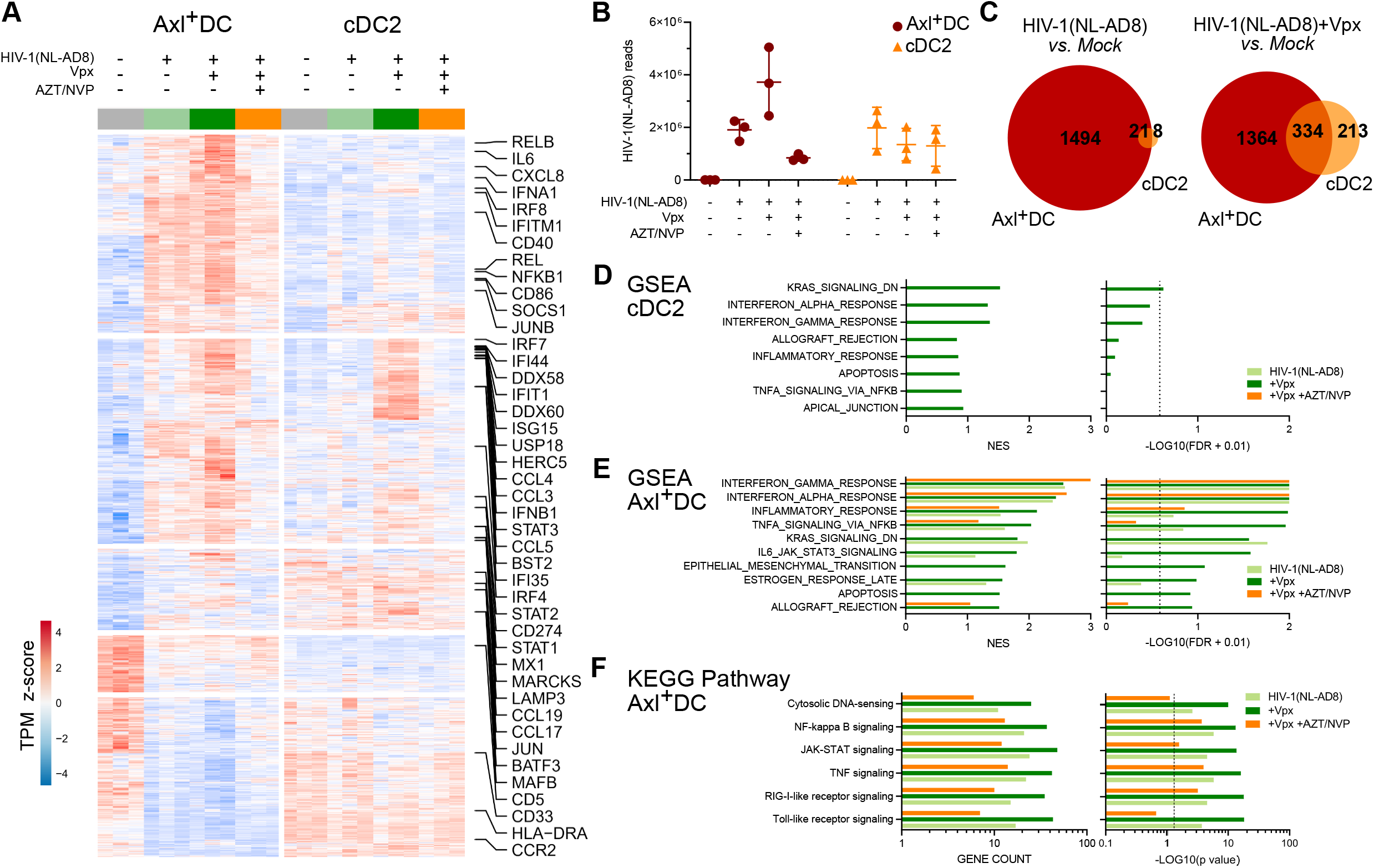
Axl^+^ DC exhibit a stronger and wider innate immune response than cDC2 after HIV-1 exposure. Bulk RNAseq was performed on purified Axl^+^ DC (AXL^+^CD123^+^CD33^int^HLA-DR^+^Lin^neg^) and cDC2 (CD1c^+^CD33^+^HLA-DR^+^Lin^neg^) infected or not for 24 h with HIV-1(NL-AD8), HIV-1(NL-AD8) + Vpx, and with or without AZT/NVP treatment. **(A)** Heatmap of normalized, centered and scaled expression (z-score) of DEGs from each HIV-1 versus mock comparisons in Axl^+^ DC and cDC2. DEGs were defined as genes with ļlog2FCļ > 1 and adjusted *p*-value < 0.05 (Benjamini-Hochberg correction), n=3 independent donors. **(B)** Quantification of HIV-1 total reads among infected or non-infected Axl^+^ DC (red) and cDC2 (yellow), n= 3 independent donors. Individual donors are displayed with bars representing SD. **(C)** Venn diagrams displaying the number of upregulated DEGs obtained in Axl^+^ DC (red) or cDC2 (yellow) when comparing HIV-1 (NL-AD8) (left diagram) or HIV-1(NL-AD8) + Vpx (right diagram) versus mock. **(D)** Top Hallmarks significantly enriched in GSEA analysis performed on ranked DEGs from each HIV-1 versus mock comparisons in cDC2 and **(E)** in Axl^+^ DC. Bar plots represent normalized enriched score (NES) and log10-transformed false discovery rate (FDR). Enrichment is considered significant when FDR < 0.25 (dashed line). **(F)** Union of top 5 pathways from KEGG database significantly enriched in DEGs from 3 HIV-1-treated conditions versus mock comparisons in Axl^+^ DC. Values are reported for the 3 conditions. Bar plots represent number of genes enriched (gene count) and log10-transformed p value. Dashed line represents p value cut-off of 0.05.

### HIV-1 exposure activates two responses in Axl^+^DC, distinct from the one induced by viral replication

To analyse in Axl^+^DC the relation between the different sensors and signalling pathways activated by HIV-1 exposure, we performed single-cell RNAseq using a droplet-based single-cell 3’ assay (10x genomics). We cultured Axl^+^DC isolated from a healthy donor with or without Vpx-complemented HIV-1(NLAD8) for 12 h or 24 h, in the presence or not of AZT/NVP for 24 h. From these five samples, we obtained a total of 2983 cells after exclusion of low-quality cells (see Methods). Quantification of viral sequences in the 24-HIV sample (Axl^+^DC exposed to HIV-1 for 24h) revealed a bimodal distribution (Figure 2A) allowing to discriminate productively infected cells (i.e., cells with a viral UMI fraction > 1.58%), containing both viral mRNA and genomic RNA, from cells that only contained viral genomic RNA from up taken virions. Importantly, the amount of productively infected cells measured by scRNAseq data was consistent with the levels of *gag*-encoded protein p24^+^ Axl^+^DC measured by flow cytometry at 48 h p.i. (Figure 2B). Spliced viral sequences (Figure S2A) were exclusively detected in cells with high proportion of viral transcripts, further supporting that viral replication had taken place in these cells. The relative contribution of each HIV-1 gene to the total viral transcripts across the productively infected cells was independent of the amount of viral mRNA in the cell (Figure S2A). Our results show that cells containing incoming viral particles can be distinguished from cells experiencing active viral replication by scRNAseq.

**Figure 2:**
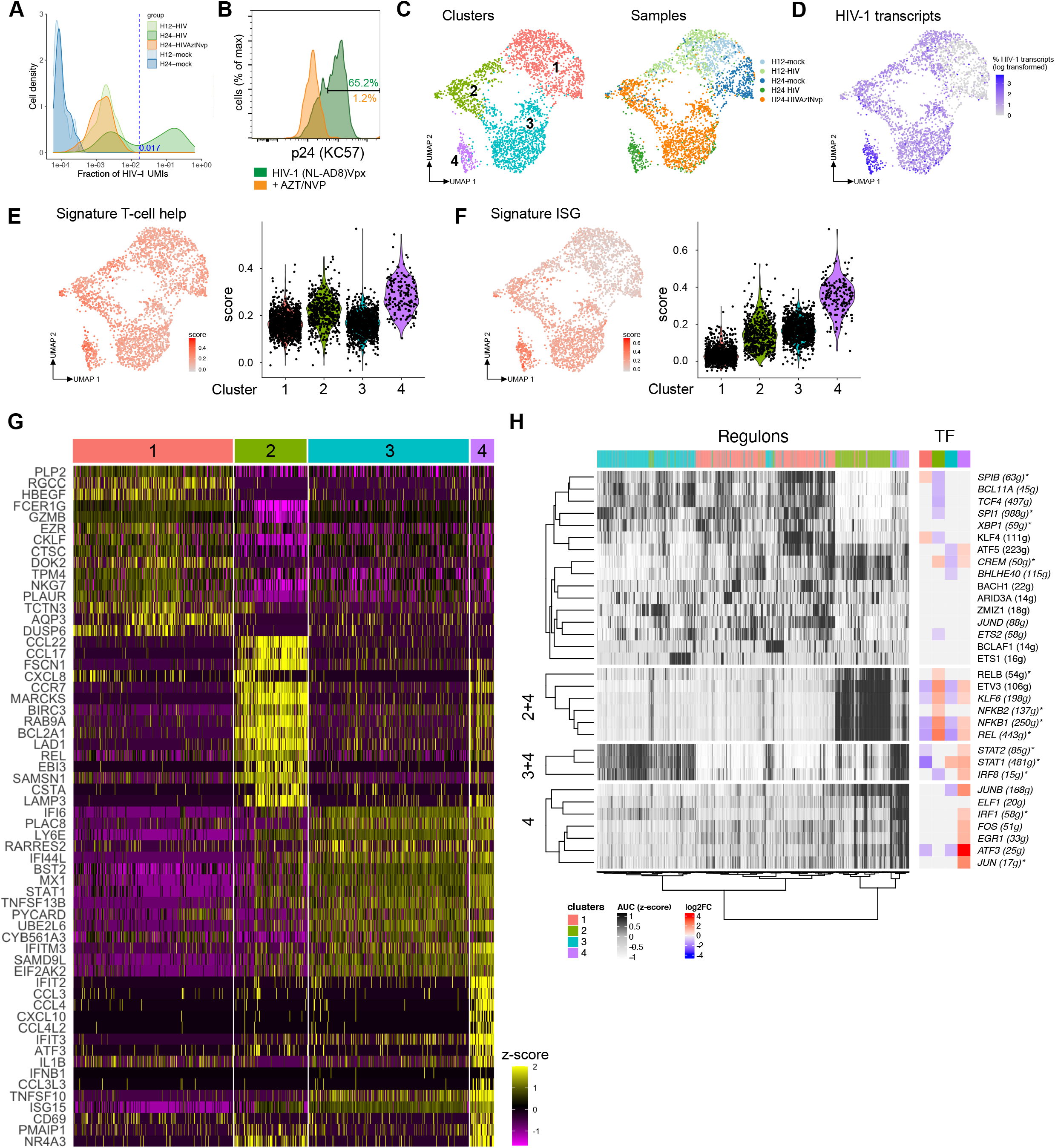
HIV-1 exposure of Axl^+^ DC induced two transcriptional responses. Purified Axl^+^ DC (AXL^+^CD123^+^CD33^int^HLA-DR^+^Lin^neg^) were cultured for 12 or 24 hours in the presence or not of HIV-1(NL-AD8), HIV-1(NL-AD8) + Vpx with or without AZT/NVP treatment before being processed for single-cell RNA sequencing. **(A)** Distribution of the HIV-1 viral UMI (unique molecular identifier) fraction computed over the total UMI counts across cells. The dashed blue line separates productively and non-productively infected cells (see Methods). **(B)** Quantification of p24 (FACS) in Axl^+^ DC infected with HIV-1(NLAD8) + Vpx (green) or HIV-1(NL-AD8) + Vpx and AZT/NVP (orange) for 48 hours. Same donor as in scRNAseq. **(C)** Uniform Manifold Approximation and Projection (UMAP) of the five pooled scRNA-seq samples, colored either by cluster (left panel) or by sample (right panel). (See methods for cluster determination). **(D)** UMAP colored by viral UMI fraction, scaled and log10-transformed. **(E-F)** UMAP plots (left) and violin plots (right) of **(E)** T cell Help Signature score and **(F)** ISG signature score in individual cells and clusters, respectively. **(G)** Heatmap of log-transformed normalized expression levels (z-score) of the top 15 significant DEGs in each cluster (see top annotation) with respect to their complementary (log2FC > 0, adj. p-value < 0.05, detected in > 50% of the cells in the cluster). DEGs found in more than one comparison are reported only once. **(H)** Gene network inference analysis. Left: heatmap of AUC (z-score) of regulon-cell pairs. Cells are annotated according the cluster they belong to (in Fig. 2C) and regulons are labelled by the corresponding TF (the number of genes is reported in parentheses). Rows and columns are clustered using Ward’s minimum variance method computed on euclidean distances. Right: heatmap of significant (adj. p-value < 0.05) log2FC values for the TFs between each cluster and its complementary. Regulons supported by high-confidence TF binding motif annotations are marked with an asterisk (see Methods) and those detected in both replicates are shown in italic (see Figure S4).

Clustering of the five pooled Axl^+^DC samples in a reduced transcriptomic space (see Methods) identified four clusters of cells (Figure 2C and S2B). A substantial fraction of Axl^+^DC cells exposed to HIV-1 for 12 h was transcriptionally similar to the population of non-exposed cells (cluster 1, Figure 2C and S2B). In contrast, all Axl^+^DC responded to HIV-1 after 24 h and distributed in three different clusters independently of the presence of RT inhibitors. Most of productively infected cells were found in cluster 4 (Figures 2D and S2C). Thus, Axl^+^DC established 3 distinct transcriptional programs in response to HIV-1 exposure, including an early response starting at 12 h (cluster 2). We ruled out that this early response was due to a higher viral uptake by Axl^+^DC since there was no clear difference in the viral transcript fraction between clusters 1 and 2 from Axl^+^DC cultured with HIV-1 for 12 h (Figure S2D). The presence of AZT/NVP did not modify the transcriptional response of HIV-1-exposed Axl^+^DC, which remained grouped in clusters 2 and 3 (Figure 2C and S2B), suggesting that Axl^+^DC can respond to incoming viral particles independently of retro-transcription. While the majority of productively infected Axl^+^DC were found in cluster 4, the remaining ones were mostly found in cluster 3, with less viral transcripts (Figure S2C and S2E), possibly reflecting a difference in the dynamics of the HIV-1 replication cycle.

Thus, the response of Axl^+^DC to HIV-1 appears heterogeneous. Viral exposure activates two distinct transcriptional programs in a RT-independent manner, whereas productive HIV-1 replication triggers another specific program.

In cluster 2, we identified markers of DC maturation (*CCR7, CD83, CD70*) and activation (*CCL17, CCL22*) (Figures 2G and S2F). The transcription factors *NFKB1, NFKB2, RELA* and *REL* were also up-regulated in cluster 2, consistently with the enrichment of genes involved in T-cell help (Figure 2E, S2F and S3A). Functional annotation of cluster 2-specific DEGs confirmed an enrichment in genes involved in the TNF/NF-κB pathway and T-cell activation process (Figures S2G). In contrast, Axl^+^DC cluster 3 exhibited an increased expression of 48 genes classified as ISG (Kane et al., 2016), including *IFI6, IFI44L, MX1, BST2* and the transcription factor *STAT1* (Figure 2G, S2F, S3B and S3C).

The response of Axl^+^DC cluster 4 was wider and stronger, encompassing the same ISG as in cluster 3 but also 53 additional ones (Figure S3B and S3C). In addition, inflammatory cytokines (*CCL4, IL1B* and *TNF*) together with genes involved in DC maturation (e.g., *CCR7* and *CD86*) were highly expressed in Axl^+^DC cluster 4. These cells also expressed the transcription factors *STAT1/2* and *IRF7,* as well as *NFKB1* (Figure 2G and S2F).

These results reveal that HIV-1 particles induce in Axl^+^DC two main responses characterized by ISG and genes involved in T cell help. Importantly, both responses took place in different cells and were RT-independent. However, both responses were activated in Axl^+^DC when they underwent active HIV-1 replication.

### Identification of key transcription factors involved in the Axl^+^DC responses to HIV-1 exposure

To determine the key transcription factors involved in the heterogeneous Axl^+^DC response to HIV-1, we performed gene network inference using SCENIC (Aibar et al., 2017). The clustering inferred from transcription factor activity was very similar to the one directly obtained from gene expression (Figures 2H), allowing us to link the markers detected above with their putative regulators. Cluster 2 displayed a high activity of both canonical (*NFBK1* and *REL* regulons) and non-canonical (*NFKB2* and *RELB* regulons) NF-κB pathways, as well as of pathways involving *ETV3* and *KLF6* (Figure 2H and S3D). Notably, the transcription factors responsible for the expression of these regulons were up-regulated in cluster 2 and, with the exception of *NFKB2* and *RELB,* in cluster 4 as well (Figure 2H). *Thus,* in cluster 2, HIV-1 entry induced the activation of NF-κB pathways in a RT-independent manner. Interestingly, the regulons activated in cluster 3 were also found in cluster 4 and covered genes regulated by STAT1, STAT2 and IRF8, with the STAT1 regulon encompassing the highest number of genes (481 genes) (Figure 2H and S3D). Additionally, Axl^+^DC cluster 4 response also encompassed FOS/JUN, IRF1, and ATF3 regulons.

Integrating the clustering and the regulon analyses we identified for each of the two transcriptional programs observed in Axl^+^DC exposed to HIV-1 the transcription factors potentially involved. The ISG response appeared driven by STAT1/2 and IRF, while the T cell help signature by NF-κB1/2.

To evaluate the impact of Vpx in Axl^+^DC responses, we performed a second scRNAseq experiment on Axl^+^DC exposed to HIV-1 (NL-AD8) in the absence or presence of Vpx for 24 h. We projected the cells to the clusters obtained for the first experiment using scMAP (Kiselev et al., 2018). Most of the cells in this second experiment could be assigned to the transcriptomes identified in the first experiment. HIV-1(Vpx) induced in Axl^+^DC the 3 different transcriptional programs that matched the ones observed in the first experiment (Figures 2 and S4A). Notably, similar results were obtained for cells exposed to HIV-1 in the absence of Vpx (Figure S4B). Gene network inference also confirmed the induction of the two different transcriptional programs previously observed, respectively involving the activation of the NF-κB pathways or of STAT1/2 and IRF, leading to ISG expression (Figure S4C-D).

### Axl^+^DC response to HIV-1 is independent of CD11c expression

Axl^+^DC have been proposed to represent a mix of cells sharing characteristics closer to cDC2 or pDC based on the presence or absence of CD11c expression (Leylek et al., 2019; Villani et al., 2017). To determine whether the dual response we observed in Axl^+^DC exposed to HIV-1 comes from this heterogeneity, Axl^+^DC were sorted into CD11c^+^ and CD11c^-^ cells, cultured for 24 h with HIV-1 and analyzed by RNAseq (Figure S5A). PCA analyses of gene expression revealed indistinguishable response to HIV-1 exposure in both cell populations (Figure S5B). The vast majority of DEGs specific for cells exposed to HIV-1 vs mock control, were shared among the two cell populations (n=746 out of 1113 and 895 DEG for CD11c^+^ and CD11c^-^ cells, respectively, Figure S5C). Further analyses of DEGs by Gene Ontology, Reactome or KEGG pathway showed that both CD11c^+^ and CD11c^-^ Axl^+^DC responded to HIV-1 by inducing the expression of genes involved in type-I IFN signalling, TLR and TNF signalling (Figure S5D, S5E and S5F) as shown above (Figure 1 and 2). Importantly, both CD11c^+^ and CD11c^-^ Axl^+^DC were susceptible to HIV-1 infection, as measured by HIV-1 p24 protein expression (Figure S5G). Thus, the dual response observed following Axl^+^DC culture with HIV-1 is unlikely to arise from differences in CD11c expression.

### HIV-1 sensing by Axl^+^DC activates the NF-κB pathway possibly via TLR triggering

TLR signalling was identified as one of the major pathways involved in the response of Axl^+^DC to HIV-1 (Figure 1F). Hence, we generated two additional samples, consisting of Axl^+^DC stimulated with a TLR9-ligand (CpG-A) for 12 and 24 h. We merged them with the five samples previously processed and found two new clusters, in addition to the four previously obtained (Figure 2C and S6A). Most cells from the two CpG-A stimulated samples clustered together (in cluster 5), except for a minor part of Axl^+^DC stimulated by CpG-A for 24 h, which showed a distinct transcriptome (cluster 6; see Figure S6B and S6C). Notably, markers of clusters 6 and 2 highly overlap (Figure S6C and S6E) and their relative expression is positively correlated (Figure S6D). The cells stimulated by CpG-A for 24 h showed a high expression of the maturation markers CCR7, CD83, CD40 and the cytokines CCL17 and CCL22, all involved in T-cell priming. Then, by computing regulon activity in CpG-A-stimulated cells, we found that NFκB-regulated genes were specifically induced in both cluster 2 and 6, and the extent of the response was highly similar between the two clusters (Figure S6F).

Thus, the transcriptional response observed in Axl^+^DC following 12 and 24 h culture with HIV-1 suggest that HIV-1 activates TLR signalling in this cell subset.

### STING activation is central for the induction of ISG in Axl^+^DC exposed to HIV-1

Consistent with our previous results (See et al., 2017), CpG-A stimulation of pDC and Axl^+^DC, raised high levels of *IFNB* transcripts in pDC but no detectable levels in Axl^+^DC, confirming the absence of contamination by pDC in our Axl^+^DC preparation, and thus validating our purification strategy (Figure 3A). In contrast, upon HIV-1 exposure Axl^+^DC expressed *IFNB* mRNA as seen by qPCR (Figure 3A) and by scRNAseq (Supplementary Table). Of note, the levels of *IFNB* mRNA induced by HIV-1 were similar between Axl^+^DC and pDC, in contrast to cDCs for which *IFNB* expression was absent (Figure 3A). We also confirmed that type I IFN plays a role in the induction of ISG following HIV-1 sensing. The use of a blocking Ab cocktail against IFNα, IFNβ and IFNαβ receptor led to a decrease in *MX1* expression induced by HIV-1, independently of viral replication (Figure 3B). The induction of *IFNB* gene expression was also prevented by the IFN blocking reagents in Axl^+^DC with active HIV-1 replication (Figure 3C). *IFNB* and ISG expression being the highest in cells with active viral replication, we evaluated the anti-viral effect of IFNβ on purified Axl^+^DC prior to HIV-1 exposure. HIV-1 replication was prevented in Axl^+^DC and cDC2 when treated with I FNβ (Figure S6G), confirming in these cells that type I IFN confers protection to HIV-1 prior to infection. Once cells are infected however, the induction of ISG does not protect the infected cells from HIV-1 replication.

**Figure 3:**
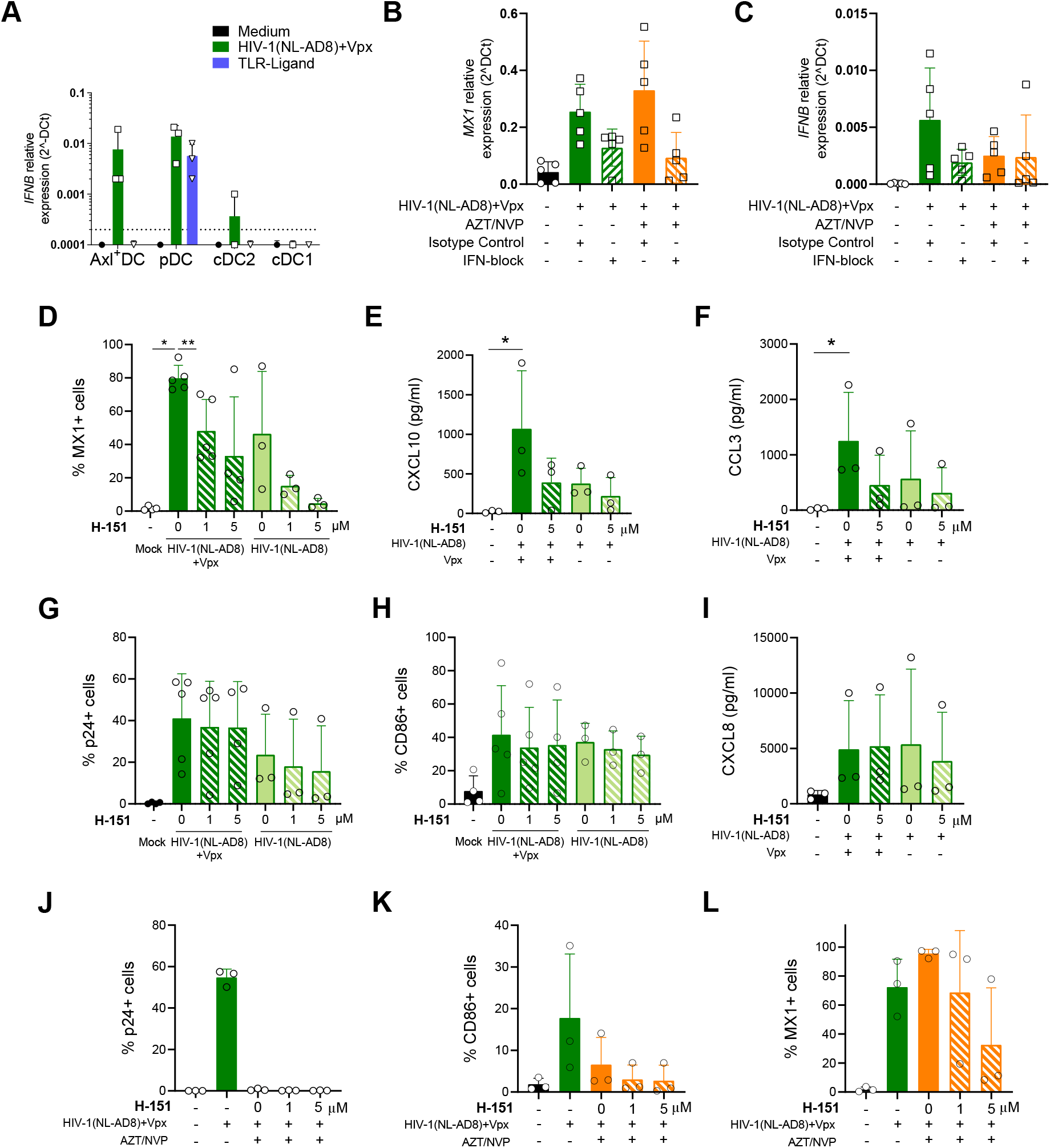
The HIV-1-induced ISG responses in Axl^+^DC are STING-dependent before and after viral retrotranscription. **(A)** *IFNB1* relative expression measured by RT-qPCR in the 4 DC subsets infected or not with HIV-1(NL-AD8) + Vpx or treated with TLR ligands (CpG-A for Axl^+^ DC and pDC, and R848 for cDC1 and CDC2), n = 3 independent donors. *MX1* **(B)** and *IFNB* **(C)** relative expression measured by RT-qPCR in Axl^+^ DC exposed or not with HIV-1(NL-AD8) + Vpx, and treated or not with AZT/NVP or with a cocktail of type I IFN blocking antibodies, n= 5 independent donors. **(D-I)** Axl^+^ DCs were treated with the STING antagonist H-151 at different doses and for 30 min before exposure to HIV-1(NL-AD8) + Vpx. After 48 hours, cell marker expression and cytokine-bead array were performed by flow cytometry. **(D)** Expression of MX1, secretion of CXCL10/IP-10 **(E)** and CCL3/MIP1#x2370; **(F)** as well as p24 **(G)**, CD86 **(H)** and the presence in culture supernatant of CXCL8/IL-8 **(I)**. n = 4 or 5 for HIV-1 (NLAD8) + Vpx samples and n= 3 for HIV-1 (NLAD8) samples. **(J-L)** Axl^+^ DCs were treated with different doses of H-151 for 30 min before exposure to HIV-1(NL-AD8) + Vpx in the presence or absence of AZT/NVP, for 48 hours. Expression of p24 **(J)**, CD86 **(K)** and MX1 **(L)** was measured by FACS, n=3 independent donors. Individual donors are displayed with bars representing SD, * *p* < 0.05, ** *p* < 0.01.

STING activation being central in MDDC response to HIV-1 (Lahaye and Manel, 2015), we tested its involvement in HIV-1 sensing by Axl^+^DC. In Axl^+^DC cultured with HIV-1 in the presence or not of Vpx, the inhibition of STING led to a decrease of MX1 expression as well as lower CXCL10 (IP-10) and CCL3 (Figure 3D, 3E and 3F). Levels of infected cells, CD86 expression and CCL8 remained unaffected by STING-inhibition (Figure 3G, 3H and 3I). MX1 reduced expression was also observed in Axl^+^DC cultured in the presence of AZT/NVP and STING inhibitors (Figure 3J, 3K and 3L). These results confirmed that Axl^+^DC sense viral incoming particles independently of retro-transcription. They also indicate that following HIV-1 sensing, STING is key in inducing ISG expression, which are further enhanced by IFNβ production.

### Priming of CD4+ T cells by HIV-1-exposed Axl^+^DC

To evaluate at the functional level the impact of HIV-1 on Axl^+^DC, we tested their capacity to prime T cells in co-culture experiments. To bypass the problem of MHC restriction (and nominal Ag) with human cells, we used the superantigen TSST-1 (toxic shock syndrome toxin) that binds to a constant region of the MHC class II β chain present on DC and can stimulate roughly 15% of human blood T cells via their TCR providing they express Vβ2 and that DC provide an efficient co-stimulatory signal (Choi et al., 1990). Comparing blood DC subsets upon addition of TSST1, Axl^+^DC were able to stimulate IL2 production by CD4^+^ T cell and their proliferation, although at slightly lower levels than cDC2 (Figure 4A, 4B, 4C and 4D). Exposure of Axl^+^DC and cDC2 to HIV-1(Vpx) enhanced the IL-2 secretion by T cells (Figure 4B) and their proliferation (Figure 4C and 4D). These results are consistent with the DC maturation program induced by HIV-1 observed at the mRNA and protein levels in Axl^+^DC (Figure 1 and S1) and suggest a role for HIV-1 replication in DC-priming of CD4^+^ T cells.

**Figure 4:**
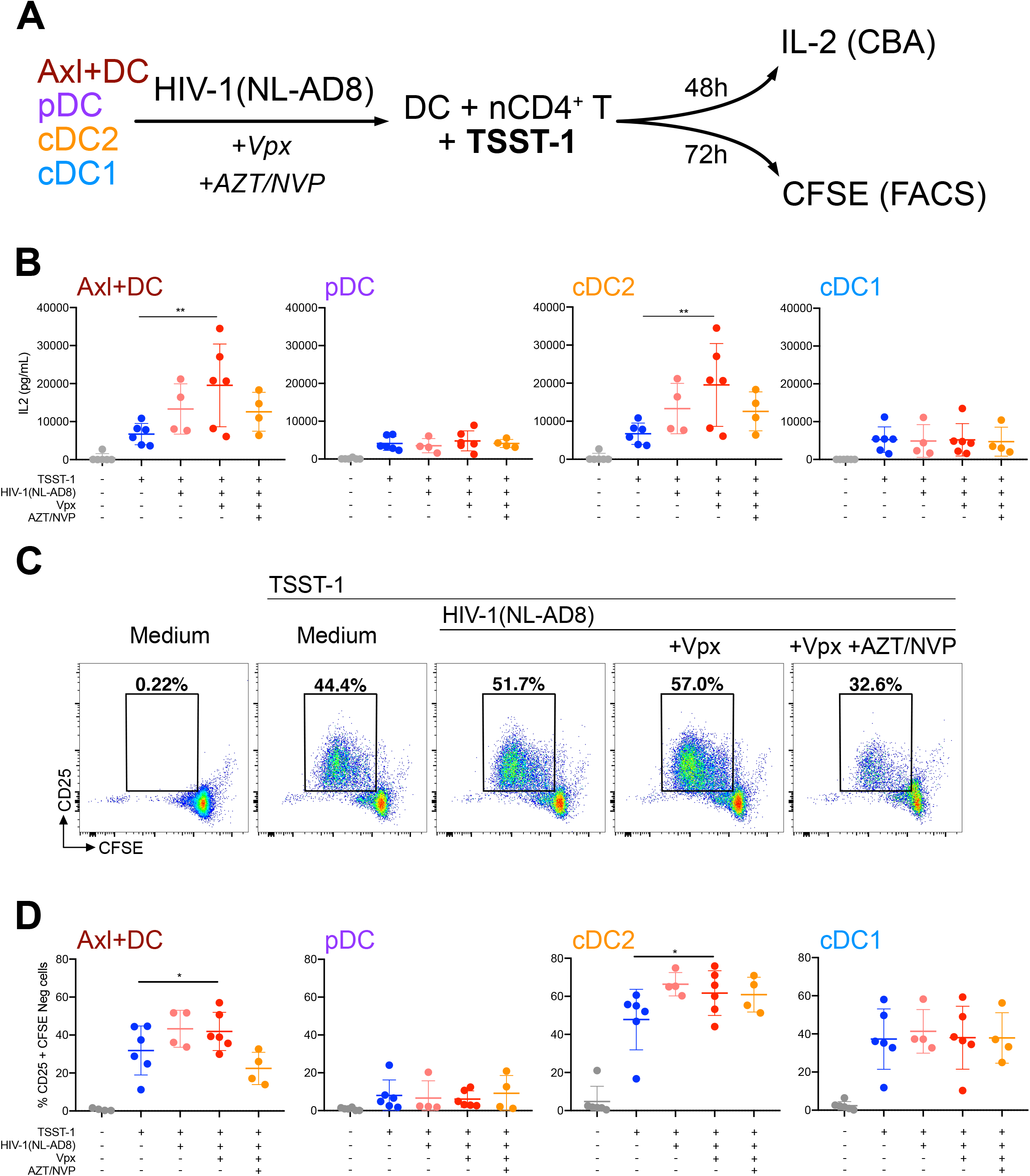
Axl^+^DC and cDC2 exposed to HIV-1 can prime naive CD4^+^ T cells. The 4 blood DC subsets were sorted and exposed to HIV-1(NL-AD8) complemented or not with Vpx and treated or not with AZT/NVP; or were stimulated with TLR-L (CpG-A for Axl^+^DC and pDC, and R848 for cDC1 and cDC2). Cells were cultured overnight, the supernatant was removed, and CFSE stained naive CD4^+^ T cells were added in a 1:10 ration (1 DC for 10 T cells) in the presence of Super Antigen TSST-1 and cultured for 48h or 72 hours for assessment of T cell proliferation. **(A)** Outline of the experiment **(B)** IL-2 quantification by CBA in the supernatant of the various co-cultures at 48 hours as indicated. n= 4 to 6 independent experiments. **(C)** Representative dotplots of CFSE and CD25 expression in gated CD3^+^ T cells following 72-hour co-culture with Axl^+^DC treated as indicated. **(D)** Quantification of proliferating CD4^+^ T cells after 72 hours of co-culture with uninfected or HIV-1-infected DC populations as in (C). Proliferating CD4 T cells were gated as CD3^+^CFSE^lo/-^CD25^+^. n= 4 to 6 independent experiments. Individual donors are displayed with bars representing SD \**p* < 0.05, ** *p* < 0.01 ***, *p* <0.001 ****, and *p* < 0.0001.

### Axl^+^DC from HIV-1 patients exhibit transcriptional programs similar to the ones induced in vitro by HIV-1

To evaluate further whether HIV-1 also impacts Axl^+^DC transcriptional program *in vivo,* we assessed DC subsets from different groups of HIV-1 infected patients. We first evaluated Axl^+^DC proportion in the blood during HIV-1 infection in a cohort of patients either during primo-infection (< less than 3 months), chronic infection or viremia suppressed with antiretroviral therapy (ART) compared to HIV-1 negative controls (Figure 5A). Frequency of Axl^+^DC among blood DC subsets remained very similar in the different groups of donors/ patients. Although we observed a tendency for pDC levels to be slightly decreased during HIV-1 primo-infection, the differences were not statistically significant. cDC2 levels also remained homogeneous, whereas cDC1 levels were significantly increased in ART-treated HIV-1-infected patients as compared to both untreated patient groups either in primo- or -chronic phase of the infection (Figure 5A).

**Figure 5:**
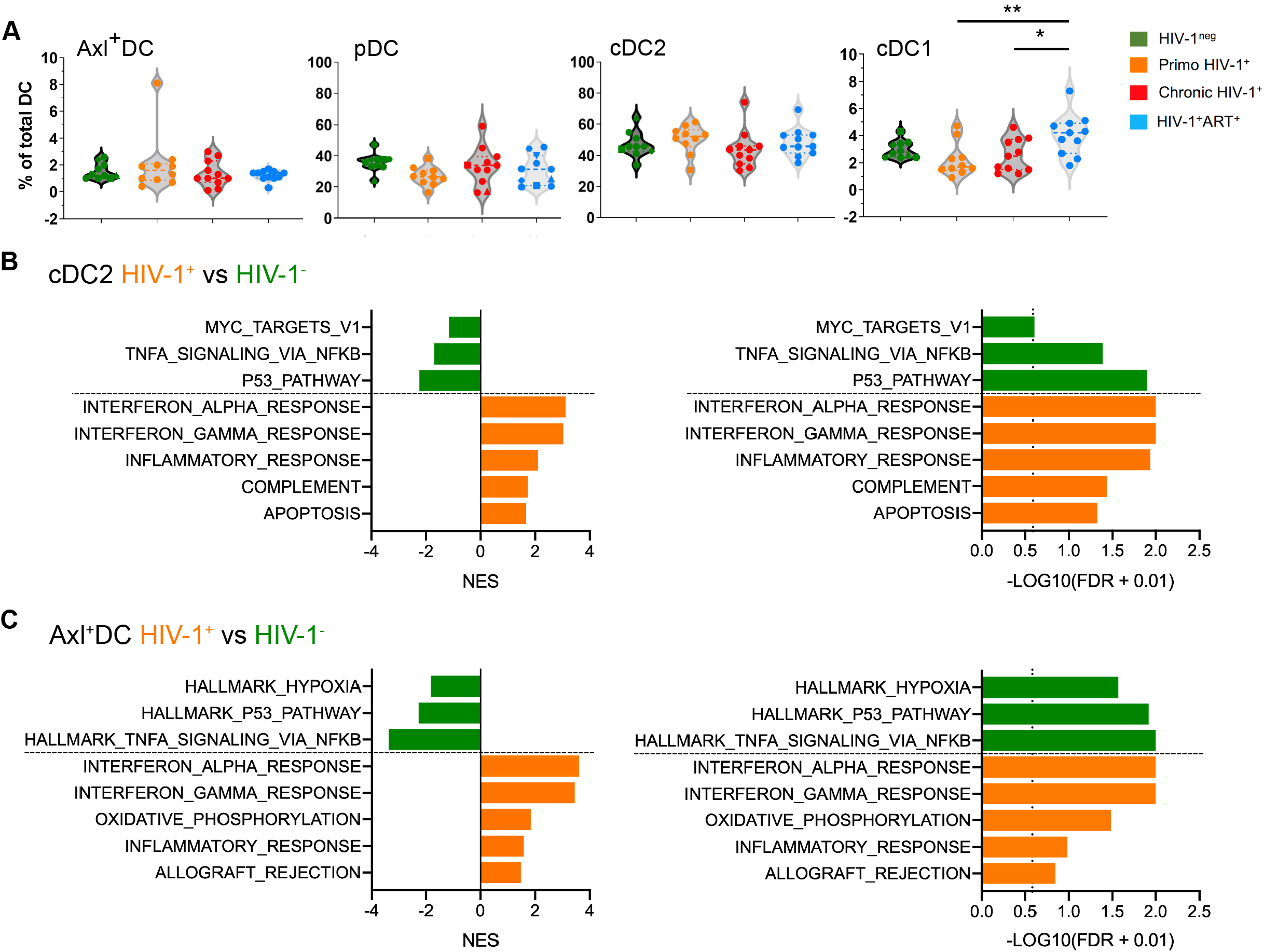
Axl^+^DC exhibit IFN and inflammatory responses in HIV-1 infected patient. **(A)** Proportion of blood Axl^+^DC, pDC, cDC2 and cDC1 in heatlhy donors or HIV-1 infected patients. **(B-C)** Hallmarks significantly enriched in GSEA analysis performed on ranked DEGs obtained comparing sorted cDC2 (B) and Axl^+^DC (C) from a HIV-1 infected patient versus DC from a healthy donor. Bar plots represent normalized enriched score (NES) and log10-transformed false discovery rate (FDR). Enrichment is considered significant when FDR < 0.25 (vertical dashed line).

We then performed scRNAseq on sorted Axl^+^DC and cDC2 populations from a HIV-1-infected patient in primo-infection and from a healthy individual as control. In contrast to *in vitro* infected Axl^+^DC, we could not detect HIV-1 transcripts in any DC population. However, functional analysis of the DEG found between DC from the HIV-1-infected patient and the control, revealed type I IFN signalling and inflammatory signatures in both Axl^+^DC and cDC2 (Figure 5B and 5C), supporting the idea that similar programs are induced by HIV-1 in Axl+DC *in vivo* and *in vitro.*

## DISCUSSION

Here, we set out to comprehensively study the response to HIV-1 of the recently identified Axl^+^DC population. We show by RNAseq analysis that primary Axl^+^DC exhibit a broader and stronger response to HIV-1 exposure compared to cDC2 that are also susceptible to the infection. Importantly, Axl^+^DC response was initiated independently from HIV-1 retrotranscription, but amplified and diversified following active viral replication, pointing to the unique sensing capacities of Axl^+^DC. HIV-1 induced functional diversity in Axl+DC as revealed by scRNAseq analysis and confirmed by functional assays. The responses induced probably originated from different mechanisms of viral sensing and depended on the amounts of viral transcripts present in each cell.

The HIV-1-induced inflammatory response we observed in a cluster (#2) of Axl^+^DC was highly similar to the one induced by TLR stimulation, suggesting that HIV-1 sensing in the endosomal pathway could be involved. Interestingly, HIV-1 replication did not take place in Axl^+^DC displaying the inflammatory response with canonical and non-canonical NF-kB pathways activated. We have previously shown that TLR stimulation switched Axl^+^DC to a HIV-1-resistant state associated with a strong inhibition of viral fusion (Ruffin et al., 2019), but further work is needed to dissect the regulation of HIV-1 fusion following its sensing by Axl^+^DC.

In contrast, the HIV-1-exposed Axl^+^DC lacking the inflammatory program expressed instead genes dependent on STAT1, STAT2 and IRF8 transcription factors, leading to the ISG response consecutive of STING activation. Although the responses were of higher intensity in Axl^+^DC with active HIV-1 replication, the ISG responses induced when RT was blocked or following active viral replication were similar, suggesting that viral RNA, proviral cDNA, or transcribed viral mRNA may constitute PAMPs eventually stimulating the STING pathway. Such responses have been previously described in MDDC (Johnson et al., 2020), but only following HIV-1 replication. Our study reveals that Axl^+^DC have the capacity to respond to HIV-1 in the absence of viral replication.

Interestingly, NF-kB and STING pathways are interconnected, since STING activation can induce the canonical NF-kB pathway via activation of the TBK1 kinase (Abe Takayuki et al., 2014). Therefore, the expression of *NFKB1* and *REL* we observed in Axl^+^DC with active viral replication could represent a consequence of STING activation. In contrast, activation of the non-canonical NF-kB pathway has been reported to inhibit STING activation (Hou et al., 2018), in line with the near complete absence of ISG response we observed in cells exhibiting a NF-kB-dependent inflammatory response. In pDC, viral exposure engaged the cells into a progressive diversification leading to 3 populations with specific responses (Alculumbre et al., 2018). In contrast, our results show the appearance of diverse responses of Axl^+^DC after viral exposure as a consequence of different mechanisms of sensing. Further work is required to evaluate the importance/extent of the NF-kB/STING signaling crosstalk in primary DC.

Using the STING inhibitor H-151, we show that STING is activated following HIV-1 sensing in Axl^+^DC, even in the presence of RT inhibitors. Thus, a cytosolic sensing of HIV-1 RNA following viral entry may take place in Axl^+^DC and induce the activation of STING and the development of a type I IFN response. Interestingly, STING can interact with RIG-1 and be activated following RIG-1 sensing of viral RNA (Solis et al., 2011), in line with our analysis of Axl^+^DC RNAseq that involved RIG-1 in HIV-1 sensing (Figure 1). Alternatively, sensing of incoming viral particles at the plasma membrane may trigger an ISG response as previously shown in macrophages (Decalf et al., 2017), via a cGAS independent STING-dependent pathway as previously proposed (Holm et al., 2016; Jakobsen et al., 2013).

Axl^+^DC specifically express Siglec-1, which endow them with a high capacity to capture HIV-1, to get infected, to produce infectious viruses in apparently internal compartments, and to transmit the virus to activated T lymphocytes (Ruffin et al., 2019). We show here that Axl^+^DC have a unique capacity to respond to HIV-1 and promote T cell activation. We reveal that the distinct sensing pathways activated by HIV-1 in Axl^+^DC instruct different outcome of the innate immune cellular response, which may impact in turn on HIV-1 viral cycle.

## AKNOWLEDGMENTS

We acknowledge the patients for their blood donation. We thank Nicolas Manel at Institut Curie for fruitful discussions, reagents, and critical reading of the manuscript. We also thank Ana-Maria Lennon-Dumenil, François-Xavier Gobert, Julie Helft, Christel Goudot, Claire Hivroz, Héloïse Delagreverie, Thimothé Bruel, Olivier Schwartz and Sebastian Amigorena for discussions or technical help. We acknowledge NGS curie, the flow cytometry facilities at Institut Curie and Institut Cochin (“plateforme Cytométrie et Immunobiologie”; CYBIO). This work was supported by grants from Agence Nationale de Recherche contre le SIDA et les hépatites virales (ANRS), Ensemble contre le SIDA (Sidaction), Laboratoire d’Excellence (Labex) DCBIOL (ANR-10-IDEX-0001-02 PSL and ANR-11-LABX-0043) to P.B., and Singapore Immunology Network (SIgN) core funding to F.G. Our work also benefitted from France-BioImaging, ANR-10-INSB-04, and CelTisPhyBio Labex (ANR-10-LBX-0038) part of the Initiative D’EXcellence (Idex) PSL (ANR-10-IDEX-0001-02 PSL). N.R. and F.B. were supported by fellowships from ANRS, Sidaction, and Fondation pour la Recherche Médicale.

## AUTHOR CONTRIBUTIONS

N.R. and P.B designed the study. N.R. and F.B performed experiments and analysed data. F.N. designed and supervised the bioinformatic analyses that were then completed by O.A.M and F.B.. C.D. and J-D.L. provided clinical samples from HIV-1 infected patients. N.R. and P.B wrote the manuscript with the help of F.B., F.N. and F.G.

---

**Supplementary Figure 1: Related to Figure 1**

**(A-B)** Venn diagrams displaying the number of upregulated DEGs (log_2_FC > 1 and adjusted *p*-value < 0.05 (Benjamini-Hochberg correction)) in Axl^+^DC **(A)** and cDC2

**(B)** obtained by comparing each HIV-1-infected sample with the corresponding mock.

**(C)** Venn diagram displaying the number of upregulated DEGs as in (A) obtained by comparing the HIV-1(NL-AD8) + Vpx + AZT/NVP condition with the mock in Axl^+^DC (red) and in cDC2 (yellow).

**(D-E)** Axl^+^DC and cDC2 were infected or not with HIV-1(NL-AD8) complemented or not with Vpx and treated or not with AZT/NVP for 48 hours. Expression of p24, MX1 and CD86 **(D)** measured by flow cytometry or quantification of CXCL10/IP-10, CCL3/MIP1#x2370; and CXCL8/IL-8 **(E)** by CBA assay. n = 3 independent donors.

**(F)** Union of Top 5 significantly enriched pathways from REACTOME database in DEGs from 3 HIV-1 exposed Axl^+^ DC conditions versus mock. Values are reported for the 3 conditions. Bar plots represent number of genes enriched (gene count) and log10-transformed p value. Dashed line represents p value cut-off of 0.05.

Individual donors are displayed with bars representing means + SD.

*p < 0.05, **p < 0.01 and ***p < 0.001

**Supplementary Figure 2: Related to Figure 2**

**(A)** Viral UMI fraction split by HIV-1 gene loci across productively infected cells (i.e., viral UMI fraction > 1.56, see Fig. 2A) in HIV-1(NL-AD8) + Vpx 24h. Cells are sorted by non-decreasing viral UMI fraction (overlayed black curve). A blue dot indicates the presence of HIV-1 RNA spliced variants in a cell.

**(B)** Cluster distribution for each sample.

**(C)** Distribution of productively infected cells across clusters (left) and their fraction within each cluster (right).

**(D-E)** Percentage of viral UMI in sample HIV-1(NL-AD8) + Vpx 12h split by cluster 1 and 2 **(D)**, or in productively infected cells from sample HIV-1(NL-AD8) + Vpx 24h split by clusters 3 and 4 **(E).**

**(F)** Volcano plots of DEGs computed between cluster pairs. ‘cl.a vs. cl.b’ indicates up-regulation (log2FC > 0) or down-regulation (log2FC < 0) in cl.a with respect to cl.b.

**(G-I)** Top 10 significantly enriched pathways from GO database (Biological Process: BP) in DEGs from clusters 2 **(G)**, 3 **(H)** and 4 **(I)** versus cluster 1 from scRNAseq analysis.

**Supplementary Figure 3: Related to Figure 2**

**(A-B)** Heatmap of log-transformed normalized expression levels (z-score) of T cell help genes **(A)** or ISGs **(B)** within DEGs from 2 by 2 cluster comparisons.

**(C)** Venn diagram displaying the number of upregulated ISGs in DEGs when comparing clusters 2, 3 and 4 with cluster 1 from the scRNAseq dataset presented in Figure 2.

**(D)** Waterfall plots of AUC values in each cell obtained from network inference analysis for the indicated regulon. Colors represent clusters from clustering analysis (5 samples, see Figure 2C) and the dashed blue lines define the AUC cutoff identifying the cells where the regulon is active (see Methods).

**Supplementary Figure 4: Axl^+^DC exposed to HIV-1 with or without Vpx complementation exhibit similar responses**

**(A-B)** Mapping of Axl+DC exposed to HIV-1(NL-AD8) +Vpx **(A)** or HIV-1(NL-AD8) **(B)** from replicate 2 (query, on the left) onto the clusters defined in replicate 1 (reference, on the right; see Figure 2). Bar sizes are proportional to the number of cells of the query that are assigned to the reference.

**(C-D)** Heatmap of AUC (z-score) of regulon-cell pairs for gene network inference analysis of the scRNA-seq dataset replicate 2. Cells are annotated according the sample they belong to (blue for mock, pink for HIV-1(NL-AD8) and green for HIV-1(NL-AD8)+Vpx) and regulons are labelled by the corresponding TF (the number of genes is reported in parentheses). Rows and columns are clustered using Ward’s minimum variance method computed on euclidean distances.

**Supplementary Figure 5: Axl^+^DC response to HIV-1 is independent of CD11c expression**

Bulk RNAseq was performed on sorted CD11c^hi^Axl^+^DC and CD11c^lo^Axl^+^DC exposed to HIV-1(NL-AD8) + Vpx or not for 24 hours.

**(A)** Representative dot plot of CD11c and AXL expression in Axl^+^DC before and after sorting cells by FACS.

**(B)** Expression of p24 measured by FACS in CD11c^+^ (red) and CD11c^-^ (blue) Axl^+^DC exposed to HIV-1 for 48 hours.

**(C)** PCA plot representation of the distribution of bulk RNAseq samples within the first 2 principal components from PCA analysis, n = 3 independent donors.

**(D)** Venn diagram displaying DEGs obtained in CD11c^hi^Axl^+^DC (red) or CD11c^lo^Axl^+^DC (blue) when comparing HIV-1(NL-AD8) + Vpx versus mock.

**(E)** Top 5 pathways from GO (Biological processes), REACTOME and KEGG databases enriched in DEGs obtained in CD11c^hi^Axl^+^DC (red) or CD11c^lo^Axl^+^DC (blue) when comparing HIV-1(NL-AD8) + Vpx versus mock. Bar plots represent log10-transformed FDR. Dashed line represents FDR cut-off of 0.05.

**Figure S6: The NFkB-dependent response to HIV-1 is largely phenocopied by TLR activation in Axl^+^ DC.**

Axl^+^ DC treated with CpG-A for 12h or 24h were processed in parallel in the experiment depicted in Figure 2 for scRNAseq analysis. The whole scRNAseq dataset considered here was therefore composed of 7 samples.

**(A)** Dimensionality reduction by UMAP of the scRNAseq dataset (7 samples). UMAP representation (UMAP1 and UMAP2 components) is colored by clusters.

**(B)** Cluster distribution within each sample.

**(C)** Heatmap of log-transformed normalized expression levels (z-score) of the top 10 significant DEGs in each cluster (see top annotation) with respect to their complementary (log2FC > 0, adj. p-value < 0.05, detected in > 50% of the cells in the cluster). DEGs found in more than one comparison are reported only once.

**(D)** Dotplot of DEGs log2FC obtained comparing clusters 2 and 6 versus cluster 1 (left panel) or comparing clusters 3 and 6 versus cluster 1 (right panel). The blue lines represent linear regressions. Pearson’s correlation coefficients and *p*-values are indicated.

**(E)** Waterfall plots of AUC values in each cell obtained from network inference analysis for the indicated regulon. Colors represent clusters from clustering analysis (7 samples, see Suppl. Figure S6A) and the dashed blue lines define the AUC cutoff identifying the cells where the regulon is active (see Methods).

**(F)** Quantification of GFP expression in blood DC subsets sorted, treated with medium or IFNb, prior to infection with HIV-1 R5GFP for 48h. n = 3 independent donors, *p<0.05 and **** p<0.0001

## MATERIAL AND METHODS

### Human subjects

Healthy individuals from Paris area donate venous blood to be used for research. Gender identity and age from anonymous healthy donors was not available. According to the 2016 activity report of EFS (French Blood Establishment), half of donors are under 40 years old, and consist of 52% females and 48% males. The use of EFS blood samples from anonymous donor was approved by the *Institut National de la Santé et de la Recherche Médicale* committee. EFS provides informed consent to blood donors. HIV-1 infected individuals (n=32) from Paris area donate venous blood to be used for research. Gender identity, age, HIV-1 viremia and CD4 T cell counts are described in Supplementary table 1. The study was approved by ethical committee (Comité de protection des personnes CPP, ID-RCB 2017-A02820-53). Written informed consent was obtained from all donors.

### Human Cell Lines

Cell lines are described in the Key Resources Table. Cell lines included 293FT, GHOSTX4R5. 293FT cells were cultured in DMEM medium, GlutaMAX (Thermo Fisher 61965-026) complemented with FBS 10% and Penicillin/Streptomycin. GHOSTX4R5 cells were cultured in DMEM medium, GlutaMAX complemented with FBS 10% and Penicillin/Streptomycin (Thermo Fisher 10378-016). All cells were cultured at 37°C with 5% CO2 atmosphere. Number of experimental replicates are indicated in the respective figure legends.

### Primary Human Cells

Peripheral blood mononuclear cells (PBMC) were isolated from buffy coats from healthy human donors (approved by the Institut National de la Santé et de la Recherche Médicale ethics committee) with Ficoll-Paque PLUS (GE). Informed consent was obtained from all donors, and samples were de-identified prior to use in the study. Total blood DCs were enriched with EasySep human pan-DC preenrichment kit (Stemcell Technologies 19251). DC enriched fraction were stained with antibodies specific for HLA-DR APCeFluor780, CD1c PerCPeFluor710 (eBioscience), CD123 Viogreen, CD45RA Vioblue (Miltenyi), Axl PE (Clone #108724, R&D Systems) CD33 PE-CF594, Clec-9A PE (BD) and with a cocktail of antibodies against lineage markers CD19 (Miltenyi), CD3, CD14, CD16 and CD34 (BD) in the FITC channel. Axl+ DC were sorted as Lin^-^ HLADR^+^ CD33^int^ CD45RA^int^ CD123^+^ Axl^+^. pDCs were sorted as Lin^-^ HLADR^+^ CD33^-^ CD45RA^+^ CD123^+^ Axl^-^. cDC2 were sorted as Lin-HLADR^+^ CD33^+^ CD45RA^-^ CD123^-^ CD1c^+^. cDC1 were sorted as Lin^-^ HLADR^+^ CD33^+^ CD45RA^-^ CD123^-^ Clec9A^+^. For the isolation of CD11^hI^ and CD11c^low^ Axl^+^ DC, CD11c APC was added to the antibody cocktail prior to FACS-sorting. All cells were sorted on a FACS Aria (BD) using Diva software (BD) in 5mL polypropylene roundbottom tubes containing 1mL X-VIVO-15 media (Lonza BE04-418F). Post-sort cell purity after gating on live cells by FCS/SSC was routinely between 90 and 99%. DCs were cultured in X-VIVO-15 complemented with Penicillin/Streptomycin.

Naïve CD4^+^ T cells were isolated from PBMC by negative selection using naïve CD4^+^ T Cell Isolation Kit (Miltenyi 130-094-131) and cultured in RPMI medium 1640, GlutaMAX complemented with FBS 10%, Gentamicin and HEPES. Number of donors and experimental replicates are indicated in the respective figure legends.

### HIV-1 production and titration

Viral particles were produced by transfection of HEK293FT cells in 6-well plates with 3μg DNA and 8μL TransIT-293 (Mirus Bio) per well. For HIV-1(NL-AD8) (described in (Freed et al., 1995)), 3μg of HIV plasmid were used. For HIV-1(NL-AD8) virus containing Vpx, 0.5μg pIRES2EGFP-VPXanyVPR and 2.5μg HIV plasmid were used. 16h after transfection, media was removed, and fresh X-VIVO-15 was added. Viral supernatants were harvested 36h later, filtered at 0.45μM, aliquoted and frozen at −80°C. Viral titers were determined on GHOST X4R5 cells as described (Mörner et al., 1999).

### HIV infection of DCs and stimulations

Sorted cells were pelleted and resuspended in complete X-VIVO-15 at 0.4 10^6^ cells/mL and 50μL were seeded in round bottom 96-well plates. In some experiments STING antagonist (H-151, Probechem) was added at 20μg/mL and cells incubated for 30 min at 37°C before adding the virus. Azidothymidine (AZT; Sigma) was used at 25 μM final concentration, and nevirapin (NVP; Sigma) was used at 5 μM final concentration. For TLR ligand stimulation, CpG-A (ODN2216, Invivogen) was used at 5μg/mL, CL264 and R848 (Invivogen) at 10μg/mL. For infections, 150μl of media or dilutions of mock or viral supernatants were added. 48hr after infection, cell culture supernatants were harvested, after neutralization with 10% NP40 lysis buffer, for IP-10 (CXCL10), MIP1⍰ (CCL3) and IL-8 (CXCL8) quantification using BD CBA Flex Set Kits. Cells were fixed in 4% paraformaldehyde (PFA; Electron Microscopy Sciences) in PBS prior to analysis. Cells were stained with KC57 RD1 (Beckman Coulter 6604667), CD86 FITC (BD 555657) and pure MX1 (abcam ab95926) coupled with a goat anti-rabbit AF647 secondary antibody (Invitrogen A21245) and analyzed on a FACSVerse flow cytometer (BD). Data were analyzed using FlowJo v10 and Prism v7 for Mac (GraphPad).

For RNA sequencing, after 12h or 24h of culture, cells were harvested and processed with the Chromium Controller (For 10X Genomics single-Cell RNA sequencing) or lysed in Lysis buffer from Single Cell RNA purification Kit (NORGEN) for RNA extraction.

### Bulk RNA sequencing Analysis

Total RNA was isolated from FACS-sorted and HIV-1 infected blood Axl^+^ DC, Axl^+^ CD11c^hi^ DC, Axl^+^ CD11c^lo^ DC and cDC2 using a Single Cell RNA Purification Kit (Norgen 51800). Total RNA integrity was assessed using an Agilent Bioanalyzer (Agilent RNA 6000 pico Kit 5067-1513) and the RIN and DV200 were calculated. All RNA samples had a RIN > 7.0 (except 1 sample at RIN = 6.5). cDNA generation and amplification were performed with SMART-Seq v4 ultra Low Input RNA kit (Takara Bio 634891). Amplified cDNA was purified using AMPure XP Beads (Beckman Coulter A63881). The concentration of cDNA was determined by Qubit (Invitrogen Qubit dsDNA HS Assay kit Q322851) and LabChip electrophoresis (Perkin Elmer). Libraries were generated using the KAPA HyperPlus Kits (ROCHE) and quantified by Qubit. cDNA libraries were subjected to next generation sequencing using an Illumina NovaSeq6000 instrument. Sequencing reads were aligned to the human genome hg38 with TopHat. Differential gene expression analysis was performed with R (v 3.5.3) with the DEseq2 and the limma Package to remove batch/donor effect. Gene expression levels were analyzed on a base-2 logarithmic scale. Statistical test were performed and the p-values were corrected for multiple testing with the Benjamini Hochberg method. Genes with log2 fold change > 1 and adjusted-p value < 0.05 were considered differentially expressed genes (DEG). Heatmaps were produced using R packages pheatmap and ComplexHeatma (Gu et al., 2016) (version 2.8.0).

GSEA (Gene Set Enrichment Analysis) was performed using the GSEA Software (v4.1.0) and gene signatures from MSigDB (v7.0). GSEA has been performed with the default parameters of GSEAPreranked analysis (number of permutations = 1000 and number of minimum genes = 15). Principal component analysis was performed on the TPM transformed counts using R packages DEseq2 and ggplot. Pathway analysis was performed on DEG with either GeneOntology term (Biological Processes), KEGG or REACTOME databases.

### Bulk RNA-seq HIV-1 transcripts alignment + CD11c

100 bp Paired-end RNA-Seq reads were first mapped onto the hg38 genome using STAR (Dobin et al., 2013) version 2.7.3a, then the fraction of unmapped reads was mapped on the plasmid sequence (HIV-1 genome sequence (NC_001802.1) containing the HIV-2 protein sequence vpx) using the STAR version cited above. Chimeric reads were removed after each mapping. Expression levels of individual genes were obtained using featureCounts from Subread (Liao et al., 2013) version 1.5.1 and Transcripts per Million (TPM) were calculated for each gene. Differential expression analysis was performed using DESeq2 version 1.19.37 (Love et al., 2014) on raw read counts to obtain normalized fold changes (FC) and *Padj-values* for each gene. The analyses scripts are deposited in Github.

### Single-cell RNA-Seq library preparation and sequencing

We performed two distinct scRNAseq analysis, replicate 1 and 2 respectively. Axl+ DC samples infected for 12 or 24h with HIV-1 (NL-AD8)+Vpx with or without AZT treatment, or stimulated for 12 or 24h with CpG-A (TLR9 ligand). For replicate 2, Axl+ DC samples infected for 24h with HIV-1(NL-AD8) or HIV-1(NL-AD8)+Vpx.

Cellular suspensions of uninfected, HIV-1 infected or TLR9 stimulated Axl+ DC were loaded on a 10 X Chromium controller (10X Genomix) according to manufacturer’s protocol. Single-cell RNA-Seq libraries were prepared using Chromium Single Cell 3’ v2 Reagent Kit (10X Genomics) according to the manufacturer’s protocol. Briefly, after performing an emulsion, single cells were trapped into single droplets containing gel beads coated with unique primers bearing 10X barcodes, unique molecular identifier (UMI) and poly(dT) sequences. Reverse transcription reactions were performed in single droplets to produce full-length cDNA. After disruption of emulsion and cDNA clean-up, pooled cDNA were amplified by PCR for 14 cycles to obtain sufficient material used for library preparation using the Chromium Single Cell 3’ v2 reagents. Library quantification and quality assessment was performed by LabChip. Indexed libraries were subjected to next generation sequencing using an Illumina HiSeq2500 instrument.

### Single-Cell RNA-seq analysis – Read alignment and cell calling

Fastq files are obtained from BCF files using mkfastq command from CellRanger v2.1.0. A custom reference genome is created by concatenating the human reference genome GRCh38 (accession: GCA_000001405.25) and the plasmid sequence (pNL4.3 AD8). The annotation from Ensembl v91 is used for GRCh38. To annotate the plasmid genome, we use the annotation of the HIV-1 reference genome (accession: NC_001802) and mapped gene loci to the plasmid sequence. The final reference is built from protein-coding genes using cellranger mkgtf and mkref. Reads are aligned to the reference using cellranger count.

### Single-Cell RNAseq analysis – Cell filtering, clustering, and differential expression analysis

For replicate 1, filtered gene-cell expression matrices returned by cellranger count are concatenated to create two merged matrices, respectively containing five (mock and HIV-1 infected) or seven samples (mock, HIV-1 infected, or CpGA exposed), using Seurat v4.0.5. Cells with less than 200 detected genes (UMI count > 0) are filtered out.

For replicate 2, Axl+ DC are extracted from a DC mock sample by computing the highly variable genes (as above), then the top 20 PCs, and finally the clustering with parameters k=30 and r=0.1, which results in two major clusters. The Axl+ DC population is identified by matching expressed markers (AXL+, CD5+, and CX3CR1+). The gene-cell matrix for the three Axl+ samples (mock, HIV-1(NL-AD8), and HIV-1 (NL-AD8)+Vpx) are concatenated to obtain a merged matrix and processed as above.

The merged matrices obtained are library-normalized with the formula log((10000·c_g_+1)/c_t_), where c_g_ is the UMI count for gene g and ct is the total UMI count in the cell. Highly variable cellular genes (hvg) are computed from the normalized matrices using mean.var.plot method setting the average expression in [0.1, 8] and the scaled dispersion > 1. PCA is computed on the scaled hvg-cell matrices and the top n significant PCs (p-value ≤ 10^-5^; JackStraw method, default parameters) are selected and projected with Uniform Manifold Approximation and Projection (UMAP) using 30 neighbors and distance cut-off 0.3. Cells are clustered on the n PCs using the Waltman and van Eck algorithm (McInnes et al., 2020; Waltman and van Eck, 2013) with k=30 neighbors and resolution r. [n=22 and r=0.1 for the 5-sample matrix; n=30 and r=0.2 for the 7-sample matrix; a small cluster composed of contaminant cells (67 for the 5-sample matrix, 37 for the 7-sample matrix) was removed from downstream analysis]. Overall 2939 cells (5-sample) and 3885 cells (7-sample) were retained, respectively.

Differentially expressed genes (DEG) between clusters and between a cluster and its complementary are identified using wilcoxon method. Differentially expressed genes (DEGs) are defined as genes expressed in at least 10% of the cells in either of the two conditions with log2FC > 0.5 and adjusted p-value < 0.05 (Bonferroni correction).

Heatmaps and Violin plots were plotted using Seurat. Signature scores were computed using the Seurat “AddModuleScore” and the gene signature of interest. Feature plots were plotted using minimum and maximum cutoff values for each feature respectively quantile 3 and quantile 97.

### Single-Cell RNAseq analysis – Gene network inference

We used the R package SCENIC v1.1.1-9, which includes three main modules: GENIE3 for network inference from gene expression data, RcisTarget for identification of direct regulatory links, and AUCell for estimating regulon activity on each cell (Aibar et al., 2017).The input to GENIE3 is a gene-cell expression matrix that has been previously filtered to keep only the genes that are detected in at least 20 cells (UMI count > 0). Genes that are co-expressed with a TF are called modules. Note that GENIE3 identifies both positive and negative regulation links.

Spearman rank coefficient is computed for each putative TF-target link and only the positive correlations are retained (⍰ > 0.3). The modules are screened with RcisTarget v1.2.1 using the motif-TF annotation, retrieved from hg19-tss-centered-10kb-7species/mc9nr.feather. The output is a set of candidate target genes for each TF. Only the top scoring 10 TF are retained for each target gene. The gene sets obtained are called regulons. Of those, only the ones found in at least 50 out of 100 SCENIC runs computed on uniform random cell subsamples (50% of the cells each) are retained.

An AUCell score is computed for each pair of cell-regulon. The score is computed by first ranking all the genes in a cell according to their expression, and then evaluating the position of the genes in the regulon in the ranked list by AUC. The shape of the distribution obtained was evaluated using the AUC_exploreThresholds function from the AUCell R package to define a threshold of positive cells where the regulon is active.

SCENIC was run on the two concatenated matrices containing the five (replicate 1) or three samples (replicate 2), respectively, as in “Single-Cell RNAseq analysis – Cell filtering, clustering, and differential expression analysis”.

### Single-Cell RNAseq analysis – cell labelling

Cells for the Axl+ DC samples infected with HIV-1 are mapped onto the four clusters obtained for the first experiments using scmap v1.4.1. The top 500 significantly variable human genes are first selected with a modified version of the M3Drop method on the gene-cell count matrix for experiment 1 containing the cells belonging to the four clusters. For each cell in experiment 2, three correlation values (cosine similarity, Pearson and Spearman) are calculated with respect to each cluster centroid, on the variable genes previously selected. A cell is assigned to a cluster if at least two correlation values are > 0.7, otherwise is left unassigned. The result of the mapping is then plotted using plotly v4.8.0.

### Single-Cell RNA-seq analysis – Quantification of viral infection

The viral UMI fraction per cell is computed as f = log_10_((10000·c_v_+1)/ct), where c_v_ is the viral UMI count and ct is the total UMI count (cellular and viral). A normal mixture model for f on the sample exposed to HIV-1 (Ad8+vpx) for 24h (replicate 1) is computed, using normalmixEM (default parameters) from R package mixtools v1.1.0. Two distributions are predicted. Cells with f ≥ t are defined as productively infected, where t is the maximum between the 99^th^ percentile of the leftmost distribution and the 1^st^ percentile of the rightmost distribution.

Spliced viral transcripts are detected as follows. A read is considered as spliced if it maps with a gap around the tat/rev splicing junction (with a tolerance of 5 bp), unspliced if it maps to an exon and to the contiguous intron (for at least 3 bp), unclassified otherwise. A spliced transcript is defined as a UMI supported by at least 2 reads such that the fraction of spliced reads is ≥ 80% of all classified reads (spliced and unspliced).

### CD4^+^ T cells activation by DCs

For each experiment, DC populations and naïve CD4+ T cells were isolated from the same donor. Sorted DCs were pelleted and resuspended in X-VIVO-15 media at 0.4 10^6^ cells/mL and 50μL were seeded in round bottom 96-well plates. Dilutions of viral supernatants for MOI of 2 were added up to 150uL and incubated overnight at 37 °C. Supernatant was then removed and previously CFSE stained autologous naïve CD4^+^ T cells were added at a ratio of 1:10 (1 DC for 10 T cells) in the presence of Super Antigen TSST1 (0.125 to 0.5 ng/mL). After 48h or 72h of co-culture, supernatants were harvested and neutralized with NP40 10X and cells were fixed in 4% PFA in PBS. Cells were then stained with Live/Dead Near-IR Red stain, CD3 AF647 and CD25 PE (BD) and analyzed on a FACS Verse (BD). Supernatants were processed for IL-2 quantification using BD CBA Flex Set Kit.

Data were analyzed using FlowJo v10 and Prism v7 for Mac (GraphPad).

### Real-time quantitative PCR

4.10^5^ sorted DC were used ex-vivo or treated as described. Cells were lysed, RNA was extracted using RNeasy micro kit (Qiagen) or Single Cell RNA purification Kit (Norgen 51800) and reverse transcription was performed using a high-capacity cDNA reverse transcription kit (Applied Biosystems) by following the manufacturer’s instructions. Real-time quantitative PCR was performed using SYBR Green I Master (Roche) with the primers described in the Key resource table (Eurogentec).

The relative quantity of IFNB1 and MX1 mRNAs was calculated between IFNB1 or MX1 RNA Cp and RPS18 Cp by the 2-ΔCt method.

### Cytokine quantification assay by CBA

IP-10 (CXCL10), MIP1⍰ (CCL3), IL-8 (CXCL8) and IL-2 concentration were measured on non-diluted or 10-fold diluted supernatants using Human cytometric assay from BD according to the manufacturer’s protocol. Data was acquired on a FACSVers (BD) and analyzed in FlowJo v10 and GraphPad prism v7 for Mac.

### Statistical analysis

Flow cytometry and qPCR data were analyzed using Prism v7 for Mac (GraphPad). Friedman test was used followed by Dunn’s multiple comparisons tests and a p value lower than 0.05 was considered as significant (*p<0.05; **p<0.01; ***p<0.001 and ****p<0.0001).

## REFERENCES

Abe Takayuki, Barber Glen N., and Williams B. (2014). Cytosolic-DNA-Mediated, STING-Dependent Proinflammatory Gene Induction Necessitates Canonical NF-κB Activation through TBK1. Journal of Virology 88, 5328–5341.

Aibar, S., González-Blas, C.B., Moerman, T., Huynh-Thu, V.A., Imrichova, H., Hulselmans, G., Rambow, F., Marine, J.-C., Geurts, P., Aerts, J., et al. (2017). SCENIC: single-cell regulatory network inference and clustering. Nat Methods 14, 1083–1086.

Alcántara-Hernández, M., Leylek, R., Wagar, L.E., Engleman, E.G., Keler, T., Marinkovich, M.P., Davis, M.M., Nolan, G.P., and Idoyaga, J. (2017). High-Dimensional Phenotypic Mapping of Human Dendritic Cells Reveals Interindividual Variation and Tissue Specialization. Immunity 47, 1037–1050.e6.

Alculumbre, S.G., Saint-André, V., Di Domizio, J., Vargas, P., Sirven, P., Bost, P., Maurin, M., Maiuri, P., Wery, M., Roman, M.S., et al. (2018). Diversification of human plasmacytoid predendritic cells in response to a single stimulus. Nat Immunol 19, 63–75.

Beignon, A.-S., McKenna, K., Skoberne, M., Manches, O., DaSilva, I., Kavanagh, D.G., Larsson, M., Gorelick, R.J., Lifson, J.D., and Bhardwaj, N. (2005). Endocytosis of HIV-1 activates plasmacytoid dendritic cells via Toll-like receptor– viral RNA interactions. J Clin Invest 115, 3265–3275.

Choi, Y., Lafferty, J.A., Clements, J.R., Todd, J.K., Gelfand, E.W., Kappler, J., Marrack, P., and Kotzin, B.L. (1990). Selective expansion of T cells expressing V beta 2 in toxic shock syndrome. Journal of Experimental Medicine 172, 981–984.

Cribier, A., Descours, B., Valadão, A.L.C., Laguette, N., and Benkirane, M. (2013). Phosphorylation of SAMHD1 by Cyclin A2/CDK1 Regulates Its Restriction Activity toward HIV-1. Cell Reports 3, 1036–1043.

Decalf, J., Desdouits, M., Rodrigues, V., Gobert, F.-X., Gentili, Marques-Ladeira, S., Chamontin, C., Mougel, M., Alencar, B., Benaroch, P., et al. (2017). Sensing of HIV-1 Entry Triggers a Type I Interferon Response in Human Primary Macrophages. Journal of Virology 91, e00147–17.

Dobin, A., Davis, C.A., Schlesinger, F., Drenkow, J., Zaleski, C., Jha, S., Batut, P., Chaisson, M., and Gingeras, T.R. (2013). STAR: ultrafast universal RNA-seq aligner. Bioinformatics 29, 15–21.

Fonteneau Jean-François, Larsson Marie, Beignon Anne-Sophie, McKenna Kelli, Dasilva Ida, Amara Ali, Liu Yong-Jun, Lifson Jeffrey D., Littman Dan R., and Bhardwaj Nina (2004). Human Immunodeficiency Virus Type 1 Activates Plasmacytoid Dendritic Cells and Concomitantly Induces the Bystander Maturation of Myeloid Dendritic Cells. Journal of Virology 78, 5223–5232.

Freed, E.O., Englund, G., and Martin, M.A. (1995). Role of the basic domain of human immunodeficiency virus type 1 matrix in macrophage infection. Journal of Virology 69, 3949–3954.

Gringhuis, S.I., van der Vlist, M., van den Berg, L.M., den Dunnen, J., Litjens, M., and Geijtenbeek, T.B.H. (2010). HIV-1 exploits innate signaling by TLR8 and DC-SIGN for productive infection of dendritic cells. Nat Immunol 11, 419–426.

Gu, Z., Eils, R., and Schlesner, M. (2016). Complex heatmaps reveal patterns and correlations in multidimensional genomic data. Bioinformatics 32, 2847–2849.

Holm, C.K., Rahbek, S.H., Gad, H.H., Bak, R.O., Jakobsen, M.R., Jiang, Z., Hansen, A.L., Jensen, S.K., Sun, C., Thomsen, M.K., et al. (2016). Influenza A virus targets a cGAS-independent STING pathway that controls enveloped RNA viruses. Nat Commun 7, 10680.

Hou, Y., Liang, H., Rao, E., Zheng, W., Huang, X., Deng, L., Zhang, Y., Yu, X., Xu, M., Mauceri, H., et al. (2018). Non-canonical NF-κB Antagonizes STING Sensor-Mediated DNA Sensing in Radiotherapy. Immunity 49, 490–503.e4.

Hrecka, K., Hao, C., Gierszewska, M., Swanson, S.K., Kesik-Brodacka, M., Srivastava, S., Florens, L., Washburn, M.P., and Skowronski, J. (2011). Vpx relieves inhibition of HIV-1 infection of macrophages mediated by the SAMHD1 protein. Nature 474, 658–661.

Izquierdo-Useros, N., Lorizate, M., McLaren, P.J., Telenti, A., Kräusslich, H.-G., and Martinez-Picado, J. (2014). HIV-1 Capture and Transmission by Dendritic Cells: The Role of Viral Glycolipids and the Cellular Receptor Siglec-1. PLoS Pathog 10, e1004146.

Jakobsen, M.R., Bak, R.O., Andersen, A., Berg, R.K., Jensen, S.B., Jin, T., Laustsen, A., Hansen, K., Østergaard, L., Fitzgerald, K.A., et al. (2013). IFI16 senses DNA forms of the lentiviral replication cycle and controls HIV-1 replication. PNAS 110, E4571–E4580.

Johnson, J.S., Lucas, S.Y., Amon, L.M., Skelton, S., Nazitto, R., Carbonetti, S., Sather, D.N., Littman, D.R., and Aderem, A. (2018). Reshaping of the Dendritic Cell Chromatin Landscape and Interferon Pathways during HIV Infection. Cell Host & Microbe 23, 366–381.e9.

Johnson, J.S., De Veaux, N., Rives, A.W., Lahaye, X., Lucas, S.Y., Perot, B.P., Luka, M., Garcia-Paredes, V., Amon, L.M., Watters, A., et al. (2020). A Comprehensive Map of the Monocyte-Derived Dendritic Cell Transcriptional Network Engaged upon Innate Sensing of HIV. Cell Reports 30, 914–931.e9.

Kane, M., Zang, T.M., Rihn, S.J., Zhang, F., Kueck, T., Alim, M., Schoggins, J., Rice, C.M., Wilson, S.J., and Bieniasz, P.D. (2016). Identification of Interferon-Stimulated Genes with Antiretroviral Activity. Cell Host & Microbe 20, 392–405.

Kiselev, V.Y., Yiu, A., and Hemberg, M. (2018). scmap: projection of single-cell RNAseq data across data sets. Nat Methods 15, 359–362.

Laguette, N., Sobhian, B., Casartelli, N., Ringeard, M., Chable-Bessia, C., Ségéral, E., Yatim, A., Emiliani, S., Schwartz, O., and Benkirane, M. (2011). SAMHD1 is the dendritic- and myeloid-cell-specific HIV-1 restriction factor counteracted by Vpx. Nature 474, 654–657.

Lahaye, X., and Manel, N. (2015). Viral and cellular mechanisms of the innate immune sensing of HIV. Current Opinion in Virology 11, 55–62.

Lahaye, X., Satoh, T., Gentili, M., Cerboni, S., Conrad, C., Hurbain, I., El Marjou, A., Lacabaratz, C., Lelièvre, J.-D., and Manel, N. (2013). The Capsids of HIV-1 and HIV-2 Determine Immune Detection of the Viral cDNA by the Innate Sensor cGAS in Dendritic Cells. Immunity 39, 1132–1142.

Lahaye, X., Gentili, M., Silvin, A., Conrad, C., Picard, L., Jouve, M., Zueva, E., Maurin, M., Nadalin, F., Knott, G.J., et al. (2018). NONO Detects the Nuclear HIV Capsid to Promote cGAS-Mediated Innate Immune Activation. Cell 175, 488–501.e22.

Lahouassa, H., Daddacha, W., Hofmann, H., Ayinde, D., Logue, E.C., Dragin, L., Bloch, N., Maudet, C., Bertrand, M., Gramberg, T., et al. (2012). SAMHD1 restricts the replication of human immunodeficiency virus type 1 by depleting the intracellular pool of deoxynucleoside triphosphates. Nat Immunol 13, 223–228.

Leylek, R., Alcántara-Hernández, M., Lanzar, Z., Lüdtke, A., Perez, O.A., Reizis, B., and Idoyaga, J. (2019). Integrated Cross-Species Analysis Identifies a Conserved Transitional Dendritic Cell Population. Cell Reports 29, 3736–3750.e8.

Liao, Y., Smyth, G.K., and Shi, W. (2013). The Subread aligner: fast, accurate and scalable read mapping by seed-and-vote. Nucleic Acids Research 41, e108–e108.

Love, M.I., Huber, W., and Anders, S. (2014). Moderated estimation of fold change and dispersion for RNA-seq data with DESeq2. Genome Biol 15, 1–21.

Manel, N., Hogstad, B., Wang, Y., Levy, D.E., Unutmaz, D., and Littman, D.R. (2010). A cryptic sensor for HIV-1 activates antiviral innate immunity in dendritic cells. Nature 467, 214–217.

Martín-Moreno, A., and Muñoz-Fernández, M.A. (2019). Dendritic Cells, the Double Agent in the War Against HIV-1. Front. Immunol. 10, 2485.

McInnes, L., Healy, J., and Melville, J. (2020). UMAP: Uniform Manifold Approximation and Projection for Dimension Reduction. ArXiv:1802.03426 [Cs, Stat].

Mörner, A., Björndal, Å., Albert, J., KewalRamani, V.N., Littman, D.R., Inoue, R., Thorstensson, R., Fenyö, E.M., and Björling, E. (1999). Primary Human Immunodeficiency Virus Type 2 (HIV-2) Isolates, Like HIV-1 Isolates, Frequently Use CCR5 but Show Promiscuity in Coreceptor Usage. Journal of Virology 73, 2343–2349.

Rhodes, J.W., Tong, O., Harman, A.N., and Turville, S.G. (2019). Human Dendritic Cell Subsets, Ontogeny, and Impact on HIV Infection. Front. Immunol. 10, 1088.

Ruffin, N., Gea-Mallorquí, E., Brouiller, F., Jouve, M., Silvin, A., See, P., Dutertre, C.-A., Ginhoux, F., and Benaroch, P. (2019). Constitutive Siglec-1 expression confers susceptibility to HIV-1 infection of human dendritic cell precursors. Proc Natl Acad Sci USA 116, 21685–21693.

Sáez-Cirión, A., and Manel, N. (2018). Immune Responses to Retroviruses. Annu. Rev. Immunol. 36, 193–220.

Schlitzer, A., McGovern, N., and Ginhoux, F. (2015). Dendritic cells and monocyte-derived cells: Two complementary and integrated functional systems. Seminars in Cell & Developmental Biology 41, 9–22.

See, P., Dutertre, C.-A., Chen, J., Günther, P., McGovern, N., Irac, S.E., Gunawan, M., Beyer, M., Händler, K., Duan, K., et al. (2017). Mapping the human DC lineage through the integration of high-dimensional techniques. Science 356, eaag3009.

Silvin, A., Yu, C.I., Lahaye, X., Imperatore, F., Brault, J.-B., Cardinaud, S., Becker, C., Kwan, W.-H., Conrad, C., Maurin, M., et al. (2017). Constitutive resistance to viral infection in human CD141 ^+^ dendritic cells. Sci. Immunol. 2, eaai8071.

Solis, M., Nakhaei, P., Jalalirad, M., Lacoste, J., Douville, R., Arguello, M., Zhao, T., Laughrea, M., Wainberg, M.A., and Hiscott, J. (2011). RIG-I-Mediated Antiviral Signaling Is Inhibited in HIV-1 Infection by a Protease-Mediated Sequestration of RIG-I. Journal of Virology 85, 1224–1236.

Villani, A.-C., Satija, R., Reynolds, G., Sarkizova, S., Shekhar, K., Fletcher, J., Griesbeck, M., Butler, A., Zheng, S., Lazo, S., et al. (2017). Single-cell RNA-seq reveals new types of human blood dendritic cells, monocytes, and progenitors. 14.

Waltman, L., and van Eck, N.J. (2013). A smart local moving algorithm for large-scale modularity-based community detection. Eur. Phys. J. B 86, 471.

Yin, X., Langer, S., Zhang, Z., Herbert, K.M., Yoh, S., König, R., and Chanda, S.K. (2020). Sensor Sensibility—HIV-1 and the Innate Immune Response. Cells 9, 254.

