## Supplementary figures and images for "Single-cell RNA-Seq analysis reveals dual sensing of HIV-1 in blood Axl^+^ dendritic cells"

### Supplemental Figure 1

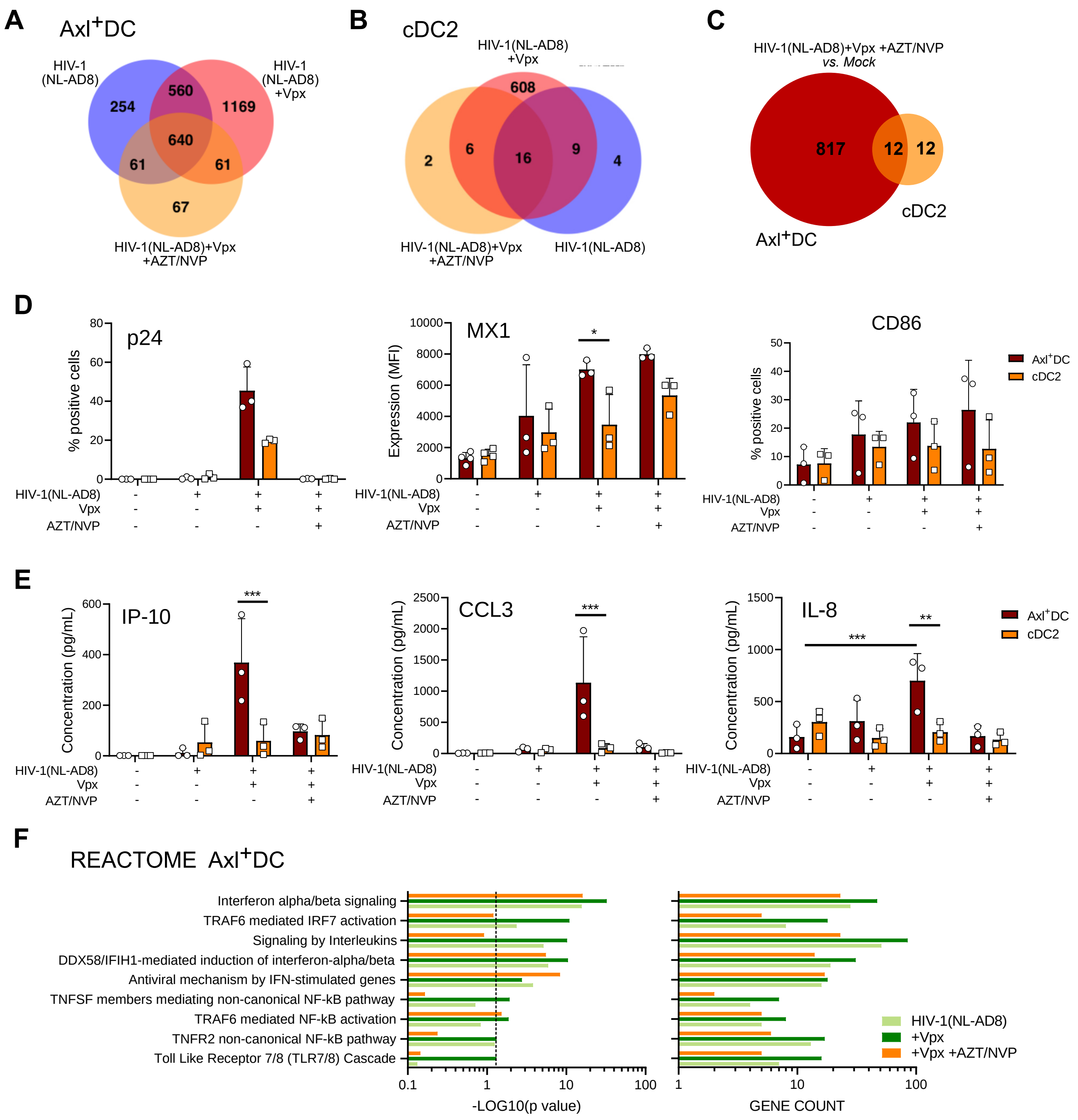

### Supplemental Figure 2

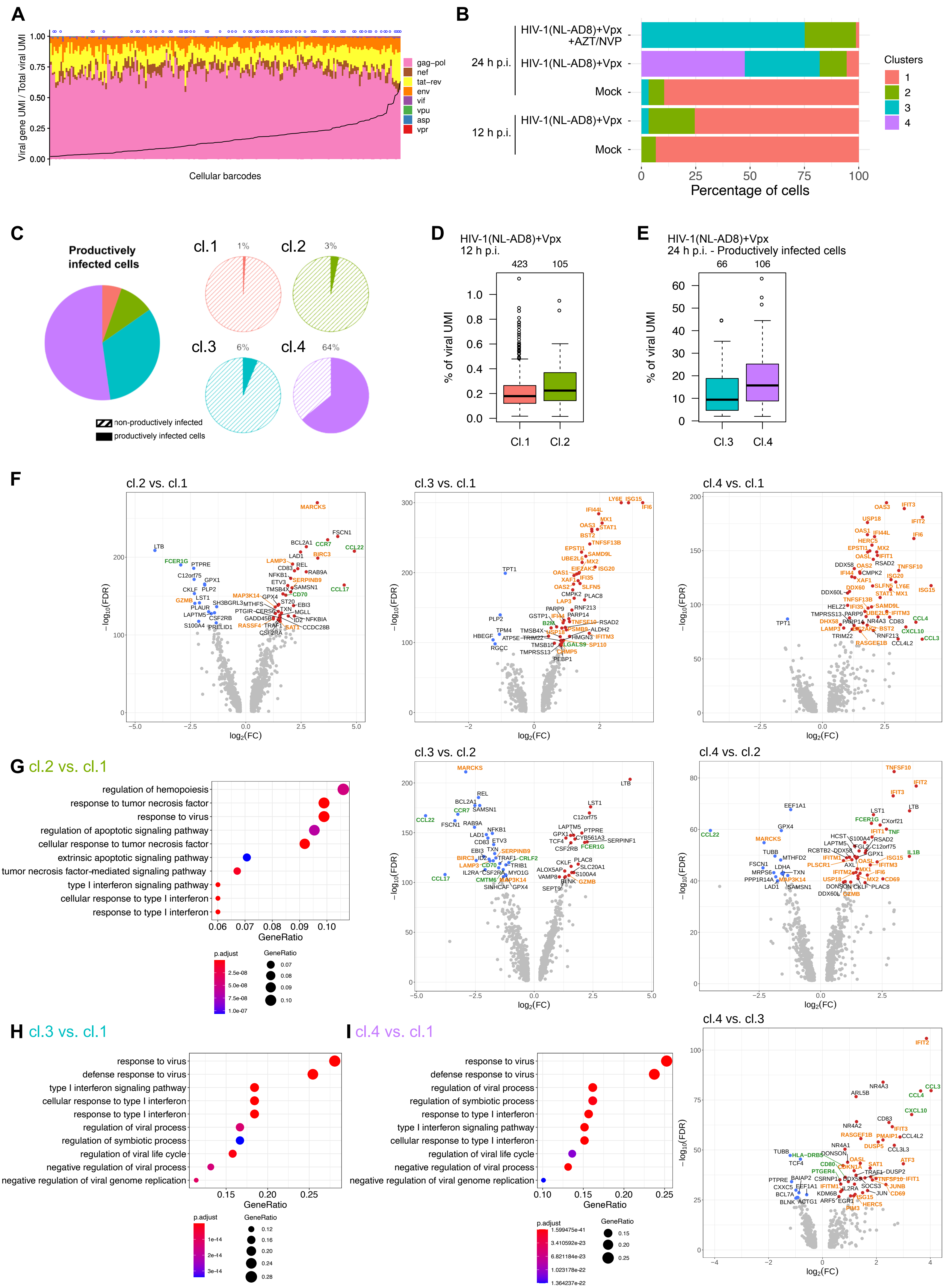

### Supplemental Figure 3

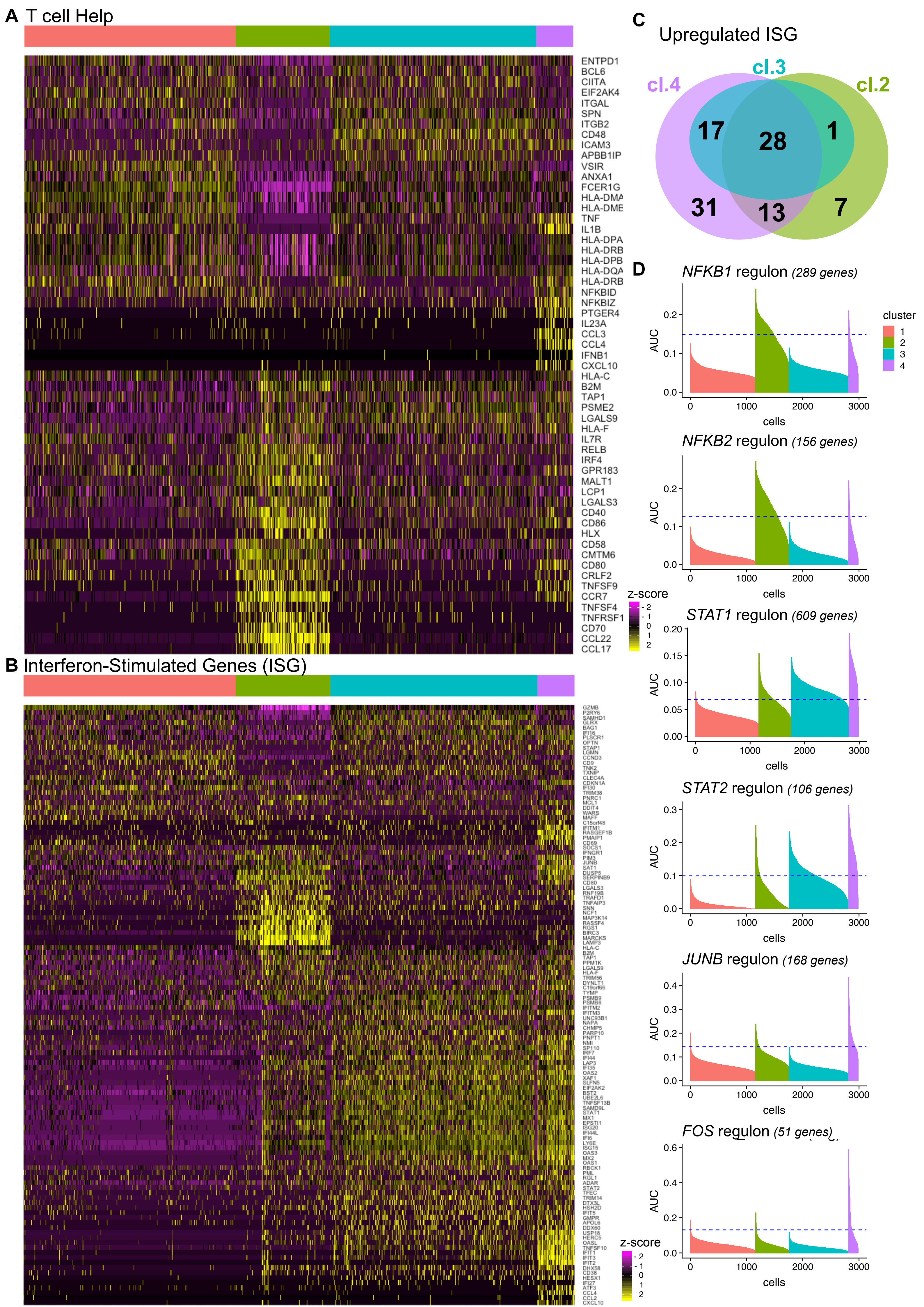

### Supplemental Figure 4

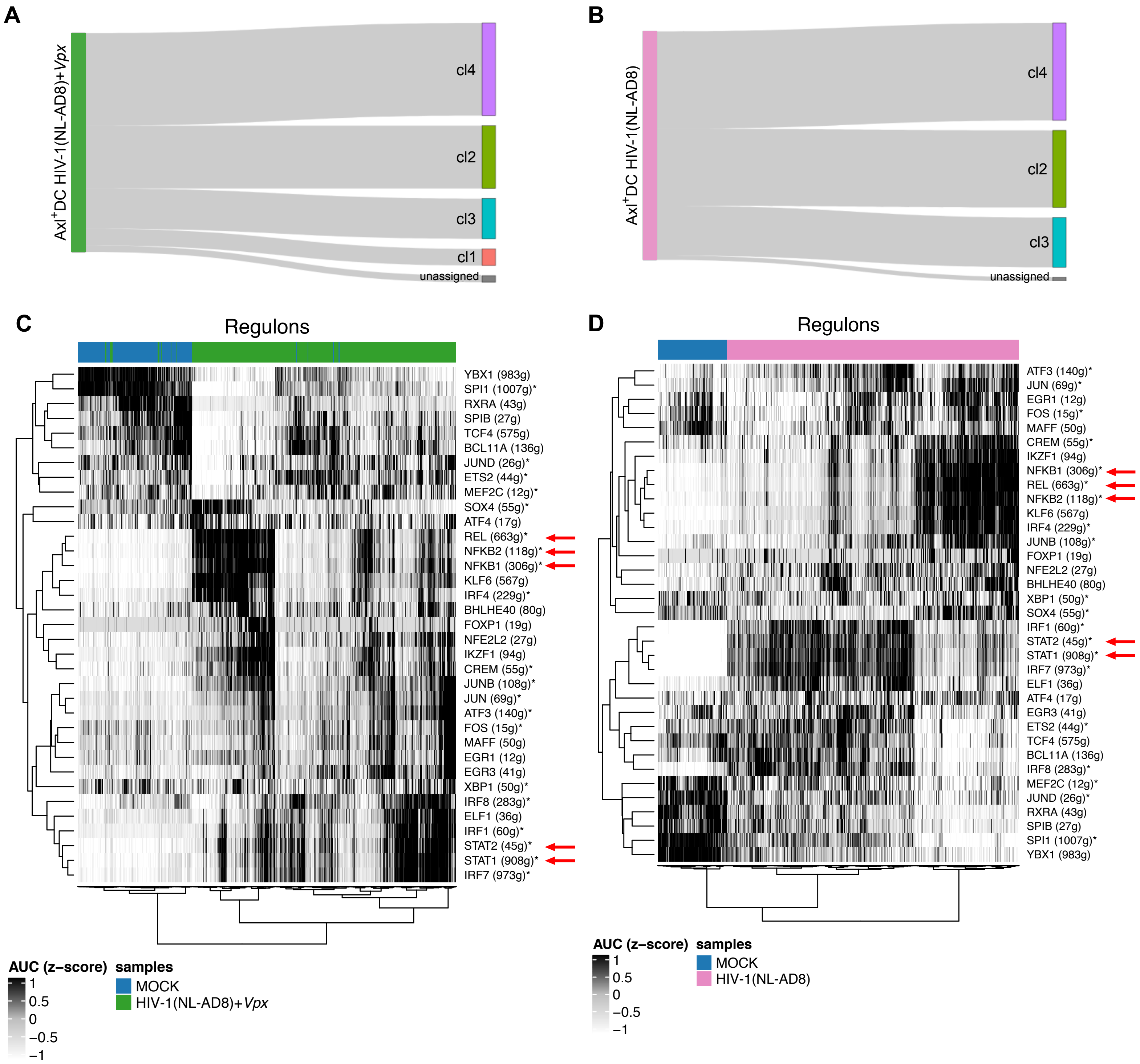

### Supplemental Figure 5

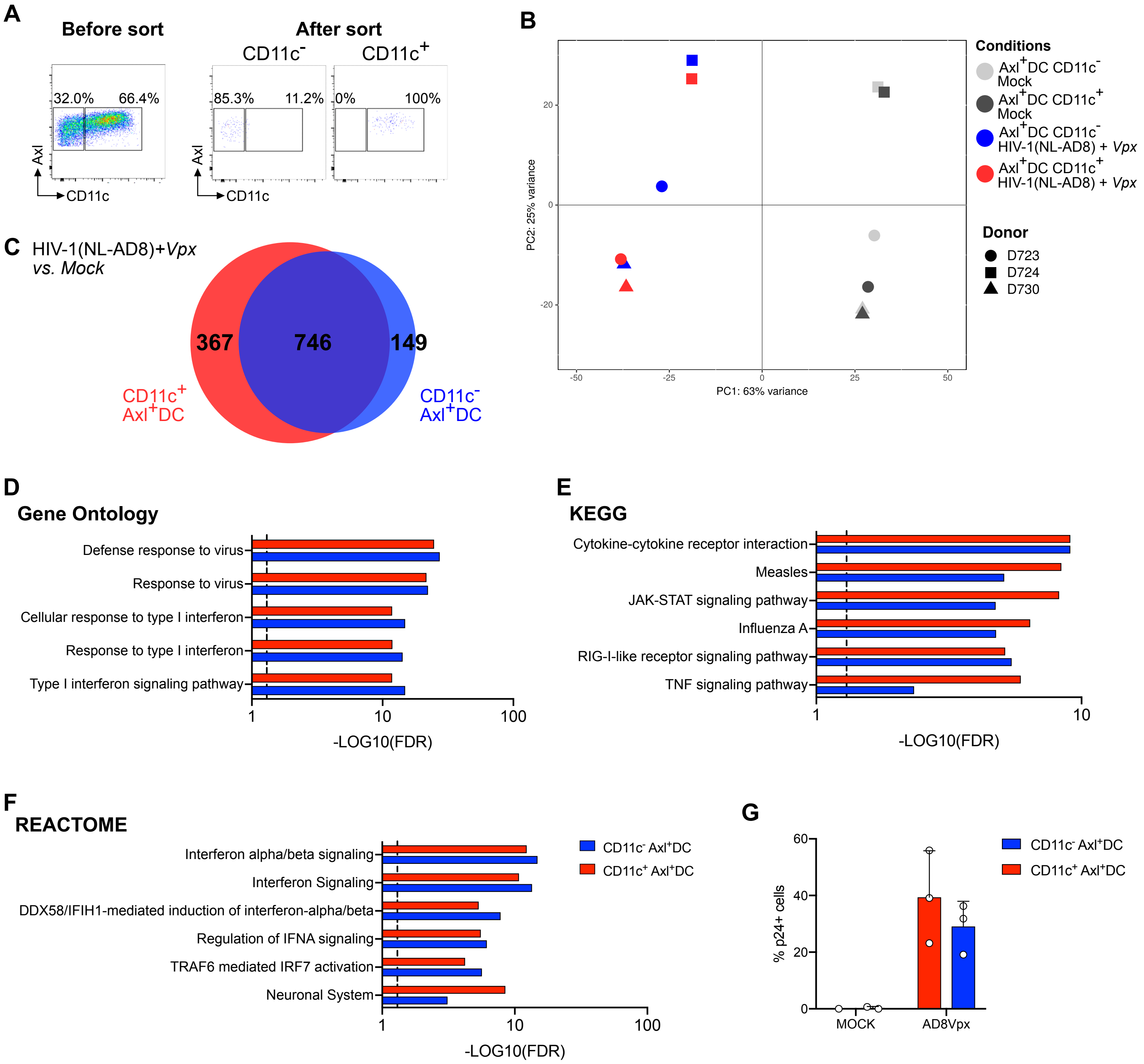

### Supplemental Figure 6

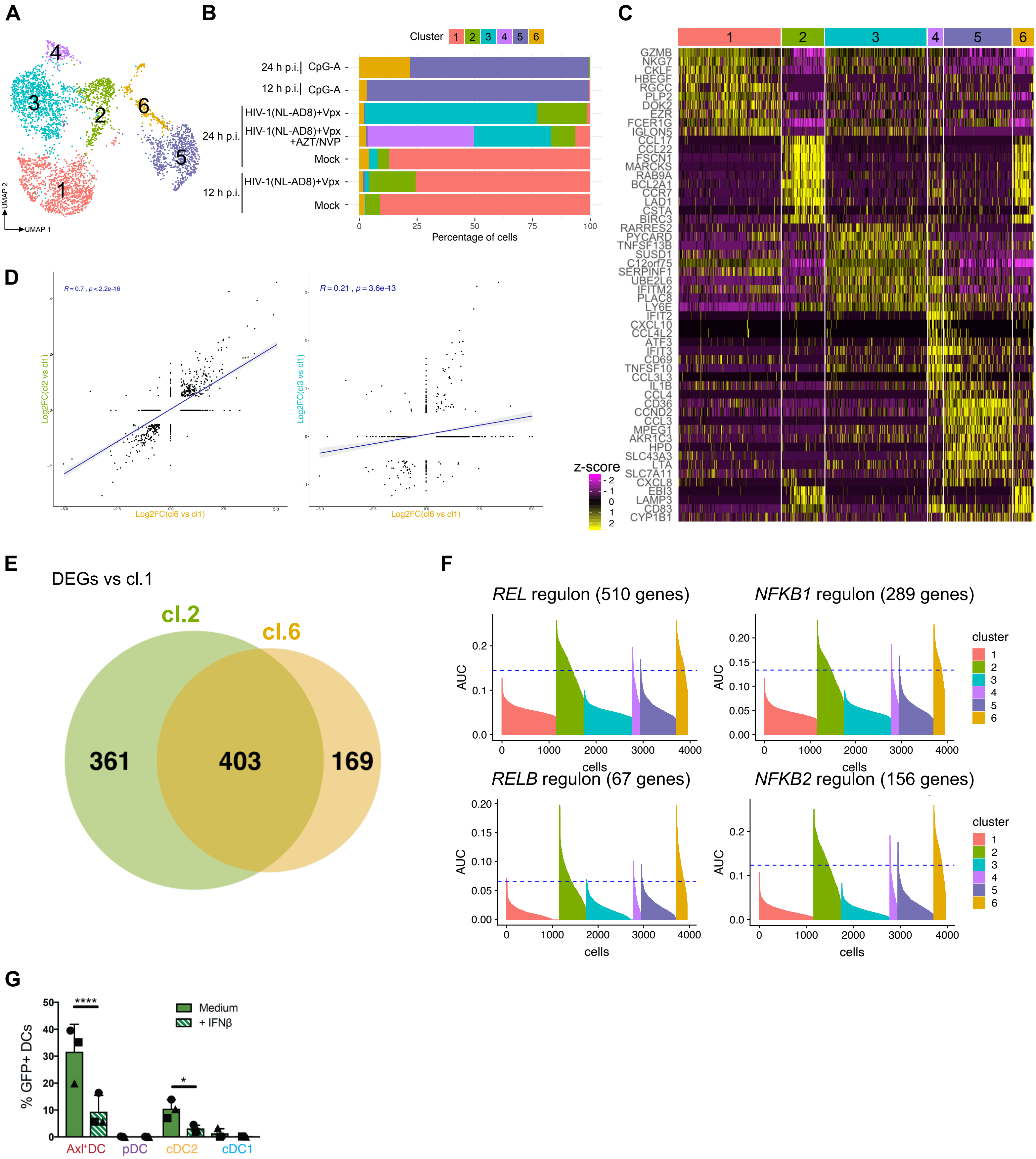
